# Allele-specific endogenous tagging and quantitative analysis of beta-catenin in colorectal cancer cells

**DOI:** 10.1101/2020.06.18.159616

**Authors:** Giulia Ambrosi, Oksana Voloshanenko, Antonia F. Eckert, Dominique Kranz, G. Ulrich Nienhaus, Michael Boutros

## Abstract

Wnt signaling plays important roles in development, homeostasis, and tumorigenesis. Mutations in β-catenin that activate Wnt signaling have been found in colorectal and hepatocellular carcinomas. However, the dynamics of wild-type and mutant forms of β-catenin are not fully understood. Here, we genome-engineered fluorescently tagged alleles of the endogenous β-catenin in a colorectal cancer cell line. Wild-type and oncogenic mutant alleles were tagged with different fluorescent proteins, enabling the analysis of both variants in the same cell. We analyzed the properties of both β-catenin alleles using immunoprecipitation, immunofluorescence and fluorescence correlation spectroscopy approaches, revealing distinctly different biophysical properties. In addition, activation of Wnt signaling by treatment with a GSK3β inhibitor or a truncating *APC* mutation modulated the wild-type allele to mimic the properties of the mutant β-catenin allele. The one-step tagging strategy demonstrates how genome engineering can be employed for the parallel functional analysis of different genetic variants.

## INTRODUCTION

Wnt signaling pathways play fundamental roles in many biological processes in development, stem cell maintenance and disease (Clevers and Nusse, 2012; Zhan et al., 2017). A key regulator of ‘canonical’ Wnt signaling is β-catenin (*CTNNB1*). When Wnt signaling is inactive, β-catenin forms part of the destruction complex, which consists of adenomatous polyposis coli (APC), axis inhibition protein 1 (Axin1) (Li et al., 2012), casein kinase 1α (CK1α) and glycogen synthase kinase 3β (GSK3β). In the destruction complex, β-catenin is N-terminally phosphorylated at Ser45 by CK1 kinase (Hagen and Vidal-Puig, 2002) and then sequentially phosphorylated at residues Ser33, Ser37 andThr41 by GSK3β (Liu et al., 2002). Subsequently, phospho-β-catenin is targeted to proteasomal degradation by the SCF^ßTrCP^ E3-ligase complex (Wu et al., 2003).

Wnt signaling is activated when Wnt ligands bind to seven-pass transmembrane receptors of the Frizzled (Fzd) protein family and the transmembrane LRP5/6 (low-density lipoprotein receptor-related protein) co-receptor (Bilic et al., 2007; Cong et al., 2004). Fzd proteins then recruit the cytosolic adaptor protein Dishevelled (Dvl) to the cell membrane *via* its DEP domain, leading to polymerization of Dvl (Schwarz-Romond et al., 2007). Dvl facilitates membrane recruitment of Axin1 by a DIX-DIX domain heterotypic interaction (Cliffe et al., 2003), which in turn stimulates GSK3β- and CK1α-mediated phosphorylation of the LRP5/6 cytosolic tail in its five PPPSPxS motifs and its binding to LRP5/6 (Stamos et al., 2014). As a consequence, the destruction complex disassembles, β-catenin is no longer degraded and translocates into the nucleus. In the nucleus, β-catenin interacts with T cell factor (TCF)/lymphoid enhancer binding factor (LEF) family of transcription factors and activates target genes in a cell-type dependent manner. In addition to its role in Wnt signaling, β-catenin is also found at cell-cell adherens junctions (AJs) mediating the interactions between the cytoplasmic domain of cadherins and the actin cytoskeleton (Aberle et al., 1994; Hoschuetzky et al., 1994; Huber and Weis, 2001).

The human β-catenin protein consists of 781 amino acids and contains three distinct structural domains. Crystal structure and NMR analysis of β-catenin reveal that both N- and C-terminal domains are structurally flexible, whereas the central armadillo repeat has a rather rigid scaffold structure (Huber et al., 1997; Orsulic and Peifer, 1996; Xing et al., 2008). The N-terminus is crucial for its stability and for cell adhesion by interacting with α-catenin. This region also contains evolutionarily conserved serine and tyrosine residues that are phosphorylated by CK1α- and GSK3β, resulting in ubiquitination and proteasomal degradation (Valenta et al., 2012). Dysregulation of Wnt signaling components is associated with a variety of human diseases, ranging from growth-related pathologies to cancer (Clevers and Nusse, 2012; Zhan et al., 2017), however, overactive Wnt signaling has been difficult to target pharmacologically (Zhong and Virshup, 2020).

Mutations in *CTNNB1*/β-catenin in colorectal cancer are typically found in its N-terminal domain, particularly in exon 3 that carries multiple phosphorylation sites for CK1 and GSK3β, including amino acids Ser33, Ser37, Thr41 and Ser45 (Cancer Genome Atlas Network, 2012; Dar et al., 2017; Jamieson et al., 2011). Mutations or deletions of these sites, often in only one allele, prevent phosphorylation of β-catenin, leading to its accumulation and subsequent activation of Wnt pathway target genes. However, the biochemical and biophysical properties of different β-catenin alleles in their endogenous loci have remained largely unknown.

To functionally analyze proteins, the introduction of small immune epitopes or fluorescent tags in-frame of the genomic locus enables their visualization and quantitative biophysical analysis. In this study, we aimed to examine the behaviour of wild-type and a mutant oncogenic β-catenin. We made use of the colon cancer cell line HCT116 which harbors one wild-type and one mutant ΔSer45 β-catenin allele to generate endogenously tagged β-catenin by CRISPR/Cas9 genome editing. We engineered a HCT116 clone with mClover-tagged wild-type β-catenin and mCherry-tagged mutant β-catenin. Tagging wild-type and mutant β-catenin in the same cell provided an opportunity to compare the function and behaviour of both proteins in parallel in the same sample. Our results indicate that wild-type and mutant β-catenin exist in two separate pools differing in their physical properties. Moreover, treatment with a GSK3β inhibitor or introduction of a truncating mutation of *APC* changed the physical properties of wild-type β-catenin, phenocopying the ΔSer45 mutant isoform.

## RESULTS

### A strategy to generate endogenously tagged fluorescent β-catenin isoforms

Previously, the dynamics and biochemical properties of β-catenin in cells were investigated by using transiently or stably overexpressed fluorescent fusion proteins (Giannini et al., 2000; Jamieson et al., 2011; Kafri et al., 2016; Krieghoff et al., 2006). To study wild-type and mutant β-catenin on a physiological level, we utilized the diploid colon cancer cell line HCT116 that harbors both a wild-type and a ΔSer45 mutant allele. This in-frame ΔSer45 deletion results in a loss of phosphorylation at this position, leading to the stabilization of β-catenin and constant activation of Wnt downstream targets (Hagen and Vidal-Puig, 2002).

To generate endogenously tagged β-catenin, we applied a gene replacement strategy using CRISPR/Cas9 editing (Mali et al., 2013; Ran et al., 2013; Yang et al., 2014) to introduce different fluorescent tags in the two alleles of the β-catenin locus of HCT116 cells in one-step (Figure 1, S1A). To this end, we identified a suitable sgRNA sequence for targeting the C-terminal region of β-catenin. This target site was chosen because it has previously been shown that C-terminal tagging did not affect its function in overexpression experiments (Giannini et al., 2000; Jamieson et al., 2011; Kafri et al., 2016; Krieghoff et al., 2006). Then, two donor templates with 180bp homology arms were designed close to the sgRNA PAM sequence (Elliott et al., 1998; Natsume et al., 2016) encoding two different fluorophores. We chose mClover3 and mCherry2 for their favorable biophysical properties such as spectral absorbance, brightness, photostability and maturation time (Bajar et al., 2016; Shen et al., 2017; Thorn, 2017). In the following, for the sake of brevity, we will refer to these two fluorescent proteins as Clover and Cherry, respectively. Moreover, FLAG and V5-tags were integrated in front of the fluorescent proteins to enable a broad spectrum of biochemical experiments. To avoid steric hindrance of the different domains, a flexible GS linker consisting of three repeats of glycine and serine residues (Gly-Gly-Gly-Gly-Ser)_n_ was inserted between the C-terminal domain of β-catenin (Trinh et al., 2004) (Figure 1, S1A). Gene resistance cassettes flanked by loxP sites were also introduced after the fluorescent proteins for selection of edited cells after homology-directed repair (HDR) (Smirnikhina et al., 2019). Selection cassettes encoded for antibiotic resistance to hygromycin and blasticidin on the Clover and Cherry donor templates, respectively (Figure S1A). The loxP sites were included to allow resistance cassette removal upon expression of Cre recombinase.

**Figure 1:**
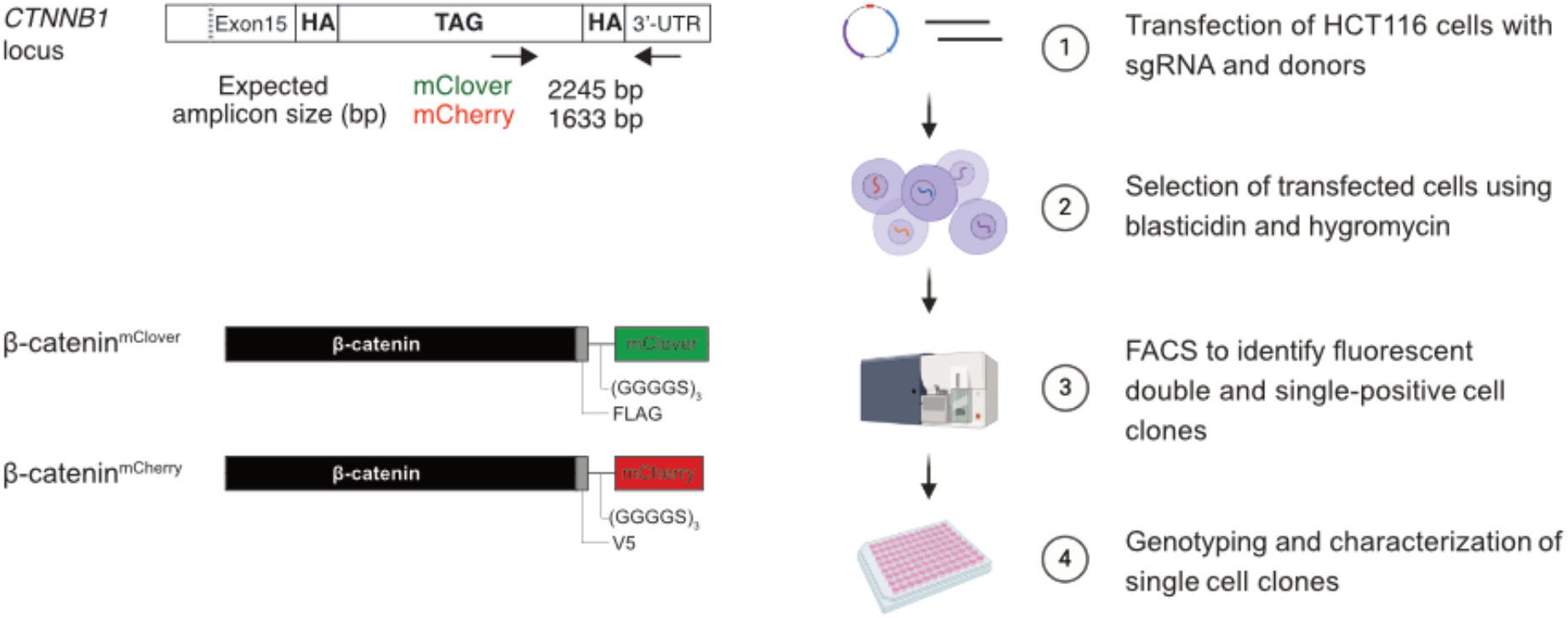
Strategy and workflow for bi-allelic fluorescent tagging of β-catenin in colon cancer cells. (Left panels) Schematic representation of *CTNNB1*/β-catenin tagging strategy, the CTNNB1/β-catenin locus and tagged β-catenin proteins. (Right panels) Workflow for the generation of endogenously tagged β-catenin in HCT116 colorectal cancer cells. See also Figure S1A for more details. HA: homology arm; UTR: untranslated region; sgRNA: short guide RNA; kDa: kilo Dalton; bp: base pair.

### Generation and characterization of β-catenin tagged cells

To generate cell lines with endogenously allele-specific tagged β-catenin, we transfected HCT116 cells with plasmids encoding for both the sgRNA targeting β-catenin (sgCTNNB1) and Cas9 protein from *S. pyrogenes* (Mali et al., 2013) and the donor template encoding either V5-mCherry BRS (blasticidinS) or FLAG-mClover HygR (hygromycin resistance) (Figure 1, S1A). Transfected HCT116 cells were first selected with puromycin for 48 h and subsequently with blasticidin/hygromycin for five days. Pooled edited cells were analyzed and sorted into single cell clones by fluorescence-activated cell sorting (FACS) (Figure 1, S1B). Infrequent fluorophore expression was observed, regardless of whether a single donor template was used (1.4% of Cherry- and 17.3% of Clover-labeled cells) or both donors were co-transfected simultaneously (0.9% of only Cherry-, 7.4% of only Clover- and 0.1% of double-labelled cells) (Figure S1B).

In total, 44 single cell clones were sorted by FACS and expanded for further characterization, which included double-positive and single-positive cells. To identify correctly edited clones, genotyping was performed using a set of primer pairs (Figure S1C). Primer pair (a) was designed to bind to the β-catenin genomic locus flanking the donor cassettes. Clone #37 showed the expected higher molecular weight bands indicating that both alleles of the *CTNNB1* were correctly edited. By contrast, clones #33, #45 and #24 showed two amplicons using primer pair (a). The lower band represents the wild-type, as observed in parental (β-catenin^wt/Δ45^) and β-catenin^wt/-^ cell lines, and the higher band represents the fluorophore-edited genomic locus. In addition, PCRs were performed using primer pairs (b) and (b’) which detect the integration of Cherry or Clover tags, respectively. As shown in Figure S1C, clones #33, #37 and #24 displayed the expected amplicon with the Clover specific primers (b’), whereas only clones #37 and #45 had the expected amplicon with the Cherry-specific primers (b). Furthermore, we used primer pair (c) binding to the homologous region of both donor cassettes and primer pair (d) binding inside the fluorescent tag. For all four clonal #33, #37, #45 and #24 cell lines, the expected bands were detected indicating correct editing (Figure S1C). In agreement with the first PCRs, clone #37 showed two bands representing correct integration of both, Clover and Cherry. As expected from the initial experiments, clones #33 and #24 showed the right molecular weight band representing Clover, whereas clone #45 showed only the lower amplicon representing Cherry. In summary, from the 44 analyzed single cell clones, only one single clone (#37) showed the correct insertion of both fluorophores in the *CTNNB1* locus (Figure S1C,D). In addition, seven clones showed a heterozygous integration of the Cherry fluorophore in the β-catenin locus and 24 clones were heterozygous for the Clover fluorophore-tagged allele. Six clones harbored an insertion in the *CTNNB1* wild-type allele presumably due to non-homologous end joining (NHEJ). Two isogenic cell lines had partial integration of the Clover tag and four clones were wild type indicating no editing events in the *CTNNB1* locus (Figure S1D). Presumably, they were selected due to random integration or high background autofluorescence of clones.

To confirm the editing events, genomic DNA, mRNA and protein lysates were analyzed and compared to the parental HCT116 (β-catenin^wt/Δ45^) cell line and to an HCT116 isogenic cell line in which the ΔSer45 allele was removed (β-catenin^wt+/-^) (Chan et al., 2002). Sanger sequencing of PCR amplicons of the target site revealed correct in-frame integration of fluorophores confirming that clone #37 harbors Clover- and Cherry-tagged β-catenin. In accordance with the genotyping results, clones #24, #33, #45 were found to be heterozygous with one remaining untagged (WT) allele. Subsequently, PCR was performed to determine which allele (WT or ΔSer45) was tagged with Clover- and which with Cherry using cDNA from reverse-transcribed mRNA of each isogenic cell line and specific reverse primers to Clover or Cherry. Sanger sequencing of cDNA templates revealed that both clones #37 and #45 carried the Cherry fluorophore in the ΔSer45 allele (Figure 2A). In contrast, #33 and #37 clones harbored the Clover fluorophore in the wild-type allele, as indicated by the in-frame sequence with Ser45.

**Figure 2:**
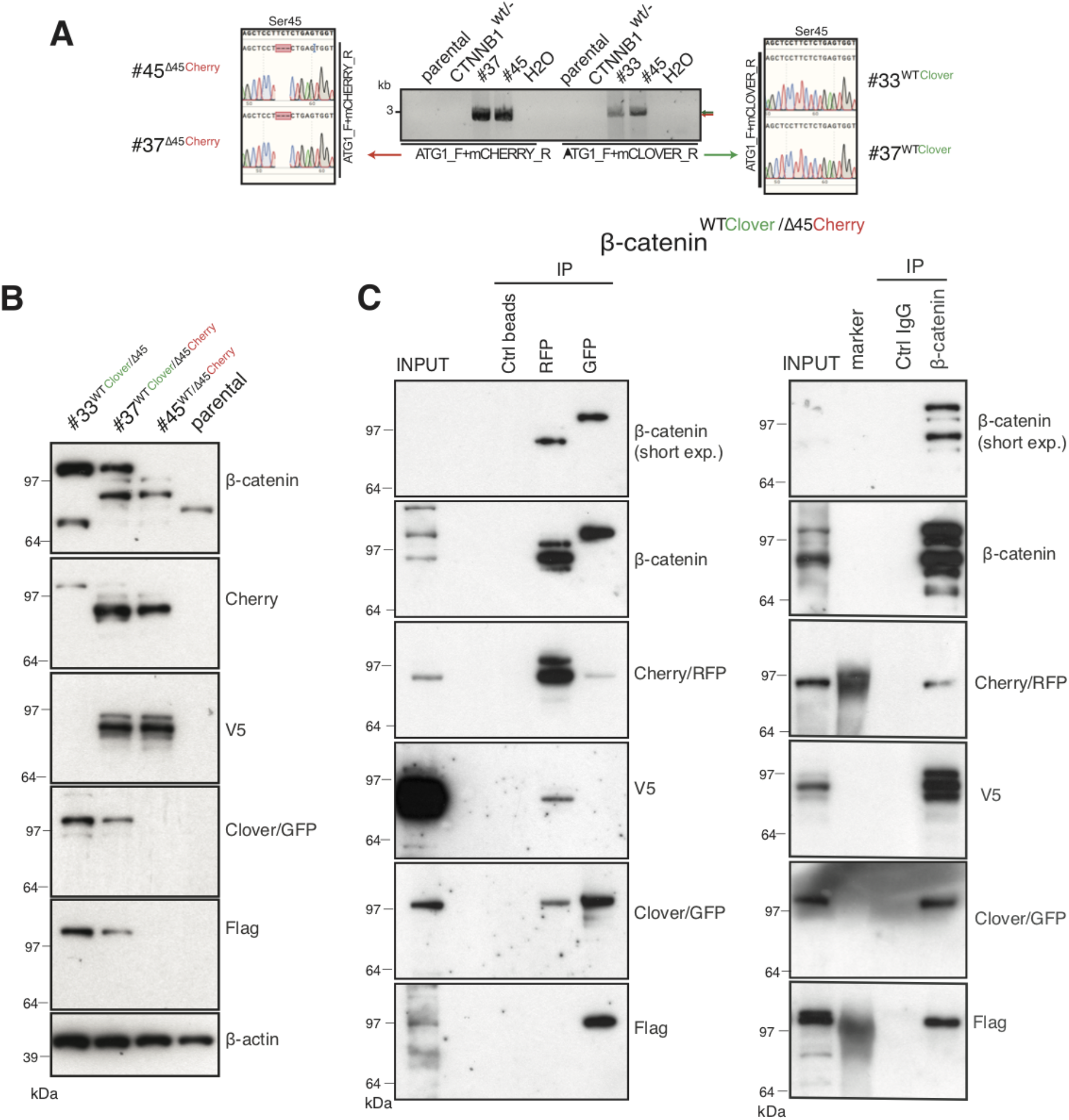
Identification and confirmation of tagged β-catenin alleles. (A) Sanger sequencing confirms bi-allelic tagging of β-catenin. Sequencing results show clones #45 (β-catenin^wt/Δ45Cherry^) and #37 (β-catenin^wtClover/Δ45Cherry^) Cherry is in-frame with the mutant allele and clones #33 (β-catenin^wtClover/Δ45^) and #37 (β-catenin^wtClover/Δ45Cherry^) Clover is in-frame with wild-type allele (codon TCT). (B) Cell lysates of indicated HCT116 cell lines analysed by western blotting with a β-catenin antibody; β-actin served as a loading control. (C) HCT116 β-catenin^wtClover/Δ45Cherry^ (clone #37) immunoprecipitation with GFP, Cherry and control beads or with a β-catenin antibody followed by immunoblotting with indicated antibodies. Representative results from three independent experiments are shown.

Next, total cell lysates of parental and isogenic knock-in cell lines were analyzed for β-catenin fusion proteins by Western blot. As shown in Figure 2B, clone #33 and clone #45 have one of the tagged variants of β-catenin, whereas clone #37 has two high molecular weight bands indicative of both tagged β-catenin variants. Furthermore, immunoprecipitations using either affinity beads for the Clover (GFP) and Cherry (RFP) fluorophores, or anti-β-catenin antibodies were performed and analyzed by Western blotting. GFP-immunoprecipitated lysates of clone #37 (Figure 2C), a β-catenin variant with apparent molecular mass higher 97 kDa was recognized by β-catenin, GFP and Flag antibodies. By contrast, upon RFP/Cherry immunoprecipitation a band around 97 kDa was detected by β-catenin, Cherry and V5 antibodies. Pulldown with a β-catenin antibody demonstrated two different β-catenin variants that were either detected by GFP/Flag or Cherry/V5 antibodies. These results confirm the presence of two differently tagged β-catenin variants in clone #37 at the protein level. Similarly, we analyzed clones #33 and clone #45 by immunoprecipitation (Figure S2). In clone #33, GFP-immunoprecipitated lysates showed a β-catenin variant higher than 97 kDa. By contrast, in clone #45 upon Cherry-immunoprecipitation β-catenin, Cherry and V5 antibodies recognized bands around 97 kDa.

Wild-type β-catenin-Clover protein runs higher than 97 kDa and it is recognized by both GFP and Flag antibodies (Figure 2B, C, S2). Mutant β-catenin-Cherry protein runs around 97 kDa as a single band, as detected in the whole cell lysate (Figure 2B, C). Upon immunoprecipitation, especially with RFP beads, several bands of mutant β-catenin-Cherry are frequently detected. To evaluate whether this is due to modifications of mutant β-catenin or due to the Cherry-tag itself, we analyzed clone #24 expressing a mutant β-catenin tagged with Clover. All three antibodies, β-catenin, GFP and Flag, detected only one band representing the mutant β-catenin-Clover protein (Figure S2), suggesting that the additional bands of mutant β-catenin-Cherry arise from the Cherry-tag. One possible explanation is that the mCherry protein undergoes chemical modification during sample preparation, for example by TCEP in the loading dye (Cloin et al., 2017). Alternatively, mCherry might undergo cleavage during chromophore maturation (Barondeau et al., 2006; Nienhaus et al., 2005; Wei et al., 2015). Importantly, such cleavage during maturation does not impair the function of the fluorophore even sometimes necessary for their functionality, as fragments remain tightly associated to form the fluorescent beta-barrel structure (Barondeau et al., 2006; Nienhaus et al., 2005).

### Functional characterization of endogenously tagged β-catenin cell lines

To test whether the C-terminal fluorescent tags of β-catenin interfere with its physiological functions, we tested the novel cell lines in different assays. First, we compared cell proliferation of the clonal cell lines to the parental β-catenin^wt/Δ45^ and the β-catenin^wt+/-^ cell line, which only had a wild-type but not a mutant β-catenin allele. All three clones #33, #37 and #45 showed a similar growth behaviour as the parental β-catenin^wt/Δ45^ cells (Figure 3A), whereas the cell line carrying a single wild-type allele (β-catenin^wt+/-^) displayed impaired growth. Next, we tested the functionality and localization of the endogenously tagged β-catenin variants. We performed siRNA-mediated silencing of β-catenin and analyzed gene expression by RT-qPCR (Figure 3B, Figure S3A). We found that downregulation of β-catenin led to the reduction in fluorophore levels as well as the Wnt target gene *AXIN2*. Consistently, depletion of β-catenin resulted in a reduction of Cherry and Clover by immunofluorescence on the protein level (Figure 3C, Figure S3A).

**Figure 3.**
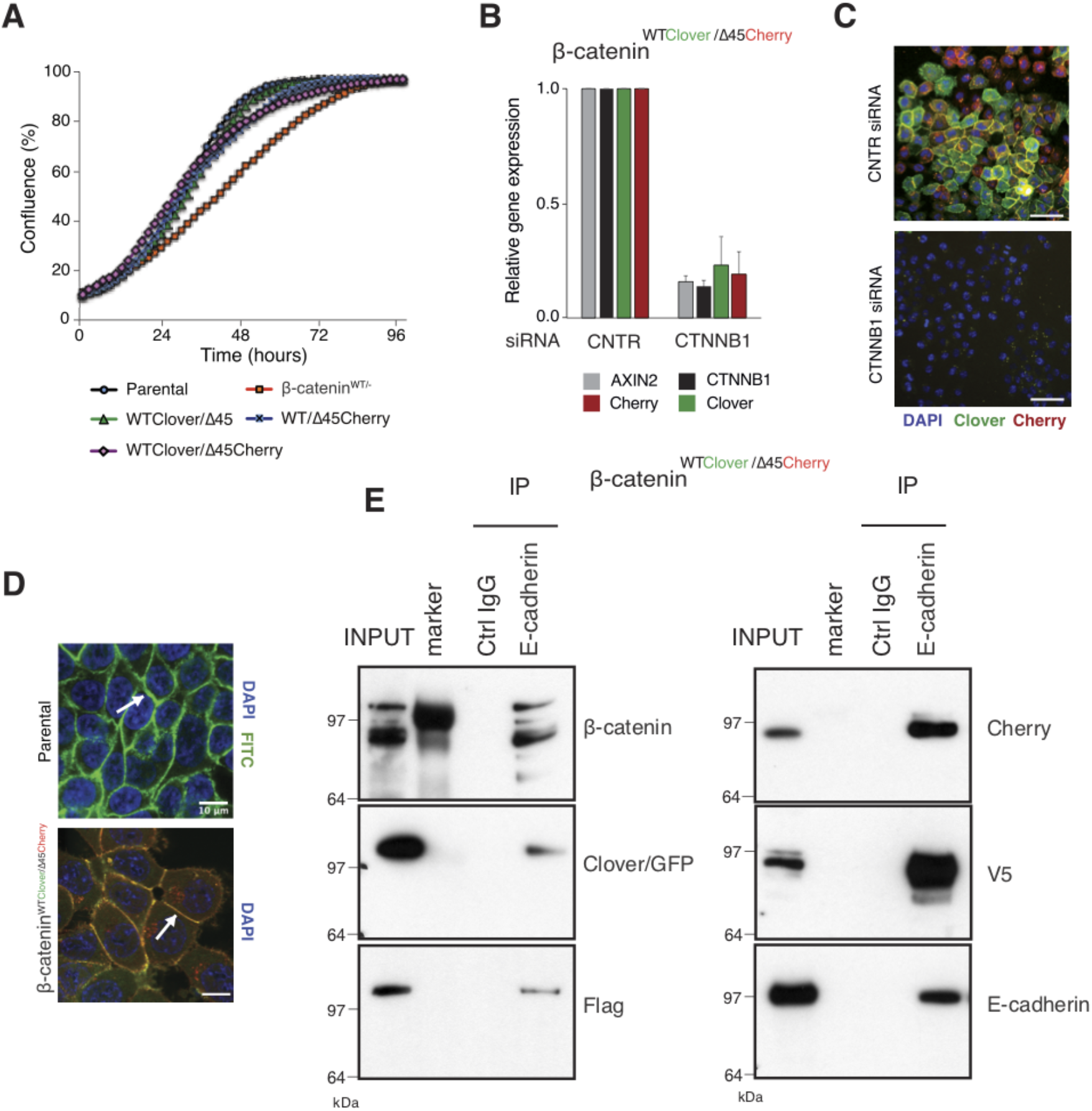
Fluorescently tagged β-catenin variants are functional and localize to adherens junctions. (A) Proliferation of the indicated HCT116 cell clones was monitored by live-cell imaging using an Incucyte instrument. (B) In HCT116 β-catenin^wtClover/Δ45Cherry^, mRNA-levels of *CTNNB1, AXIN2, Cherry* and *Clover* were determined by RT-qPCR in control conditions and upon depletion of *CTNNB1*/β-catenin by siRNA (n=5; mean ± SD). (C) Representative immunofluorescence images of HCT116 β-catenin^wtClover/Δ45Cherry^ after siRNA-mediated knockdown of *CTNNB1* is shown (n=3; scale bars: 100μm). (D) β-catenin accumulates at cell-cell junctions (arrow). Representative immunofluorescence images of HCT116 β-catenin^wtClover/Δ45Cherry^ and parental HCT116 β-catenin^wt/Δ45^ stained with a β-catenin antibody is shown (scale bars: 10μm). (E) Immunoprecipitation of HCT116 clone β-catenin^wtClover/Δ45Cherry^ with E-cadherin confirms its interaction with β-catenin. Representative results from three independent experiments are shown.

In addition to its functional role in Wnt signaling, β-catenin interacts with the cytoplasmic domains of E-cadherin and α-catenin to bridge the cell-adhesion complex and actin cytoskeleton inside the cells, thereby localizing it to the cell membrane (Aberle et al., 1994; Hoschuetzky et al., 1994). Immunofluorescence analysis confirmed the localization of the endogenously tagged β-catenin variants at the cell membrane as observed in the parental HCT116 cell line (Figure 3D, Figure S3B). Immunoprecipitation with an E-cadherin antibody demonstrated the interaction of β-catenin and E-cadherin in all clones (Figure 3E, Figure S3C). Taken together, these results indicate that endogenously tagging of β-catenin did neither affect its localization and adhesive function nor its ability to activate the Wnt pathway.

### Wild-type and mutant β-catenin variants contribute both to canonical Wnt signaling output

We next investigated the impact of endogenously tagged β-catenin on the activation of Wnt signaling. Wnt activity was measured by TCF4/Wnt-dependent reporter assay and analyzing *AXIN2* levels in the isogenic knock-in clones and parental cell line upon treatment with either the GSK3β inhibitor CHIR99021 or ICG-001, an inhibitor of CBP/β-catenin interaction (Figure 4A). Upon CHIR99021 treatment, TCF4/Wnt-reporter activity was induced and *AXIN2* gene expression levels were increased similarly in all isogenic cell lines. Treatment with the Wnt inhibitor ICG001 decreased Wnt reporter activity and *AXIN2* expression levels in all cell lines, as expected (Figure 4A). In addition, immunofluorescence analysis demonstrated that after addition of CHIR99021, nuclear β-catenin was increased in all clones indicative of increased nuclear translocation of β-catenin upon Wnt activation (Figure 4B, S4). Consistent with its mechanism of action, treatment with ICG001 did neither affect cytoplasmic nor nuclear β-catenin localization in the endogenously tagged or parental cell clones.

**Figure 4.**
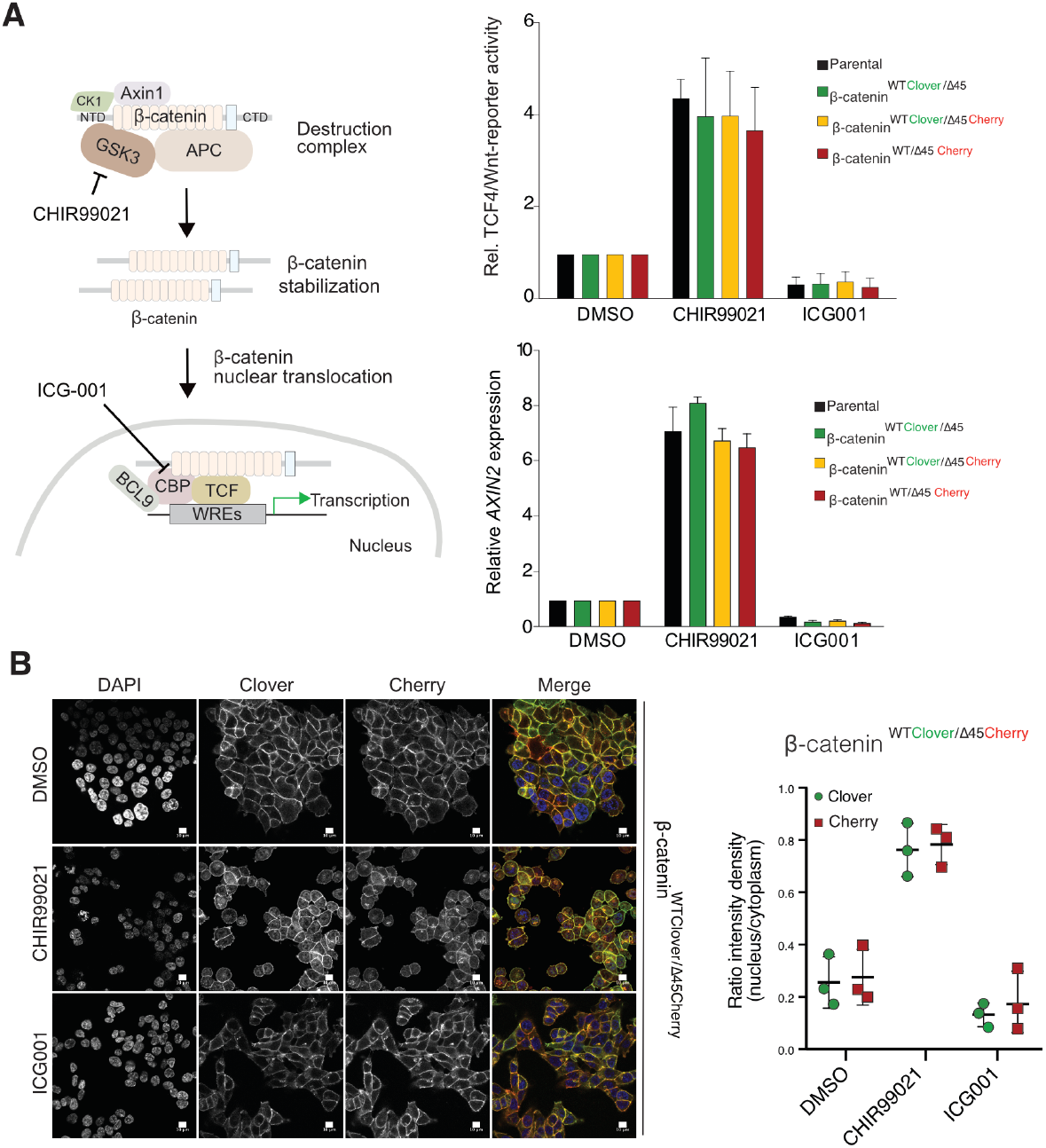
Tagging of β-catenin does not affect functionality in canonical Wnt signalling. (A) Left: Scheme showing the mode of action of GSK3β inhibitor CHIR99021 and CBP inhibitor ICG-001. Right: Indicated HCT116 cell lines were treated with 10μM CHIR99021 and 10μM ICG-001 for 24 hours, then Wnt activity was determined by a luciferase-based TCF4/Wnt-reporter assay (upper panel) and quantification of *AXIN2* mRNA-levels by RT-qPCR (n=3; mean ±SD). (B) Immunofluorescence analysis of HCT116 β-catenin^wtClover/Δ45Cherry^ after 24 hours treatment with 10μM CHIR99021 and 10μM ICG-001 is shown (scale bar: 10μm). The graph on the right depicts the ratio of nuclear to cytoplasmic fluorescent signal intensity for Clover and Cherry in HCT116 β-catenin^wtClover/Δ45Cherry^. Data from three independent experiments, each with at least 250 cells per condition, are shown as mean ±SEM. Every experiment includes at least 250 cells per condition (scale bar: 10μm). WRE: Wnt responsive element.

To analyze the impact of both wild-type and mutant variants on Wnt signaling, we performed knock-down experiments with siRNAs targeting either Cherry or Clover, thereby depleting either mutant or wild-type β-catenin alleles. Silencing of either Clover or Cherry in clone #37 reduced β-catenin and *AXIN2* expression levels in a similar manner, whereas combined knockdown of both had an additive effect comparable to siRNA β-catenin/*CTNNB1* (Figure 5A). Interestingly, silencing of wild-type β-catenin using a siRNA (GFP) resulted in a two-fold increase of mutant Cherry-tagged β-catenin that was also observed by increased immunofluorescence staining, suggesting a regulatory feedback mechanism (Figure 5B). These results indicate that both wild-type and mutant β-catenin alleles contribute to Wnt activity. This is also consistent with previous results in HCT116 cells demonstrating that Wnt secretion is essential for maintaining Wnt activity (Voloshanenko et al., 2013).

**Figure 5.**
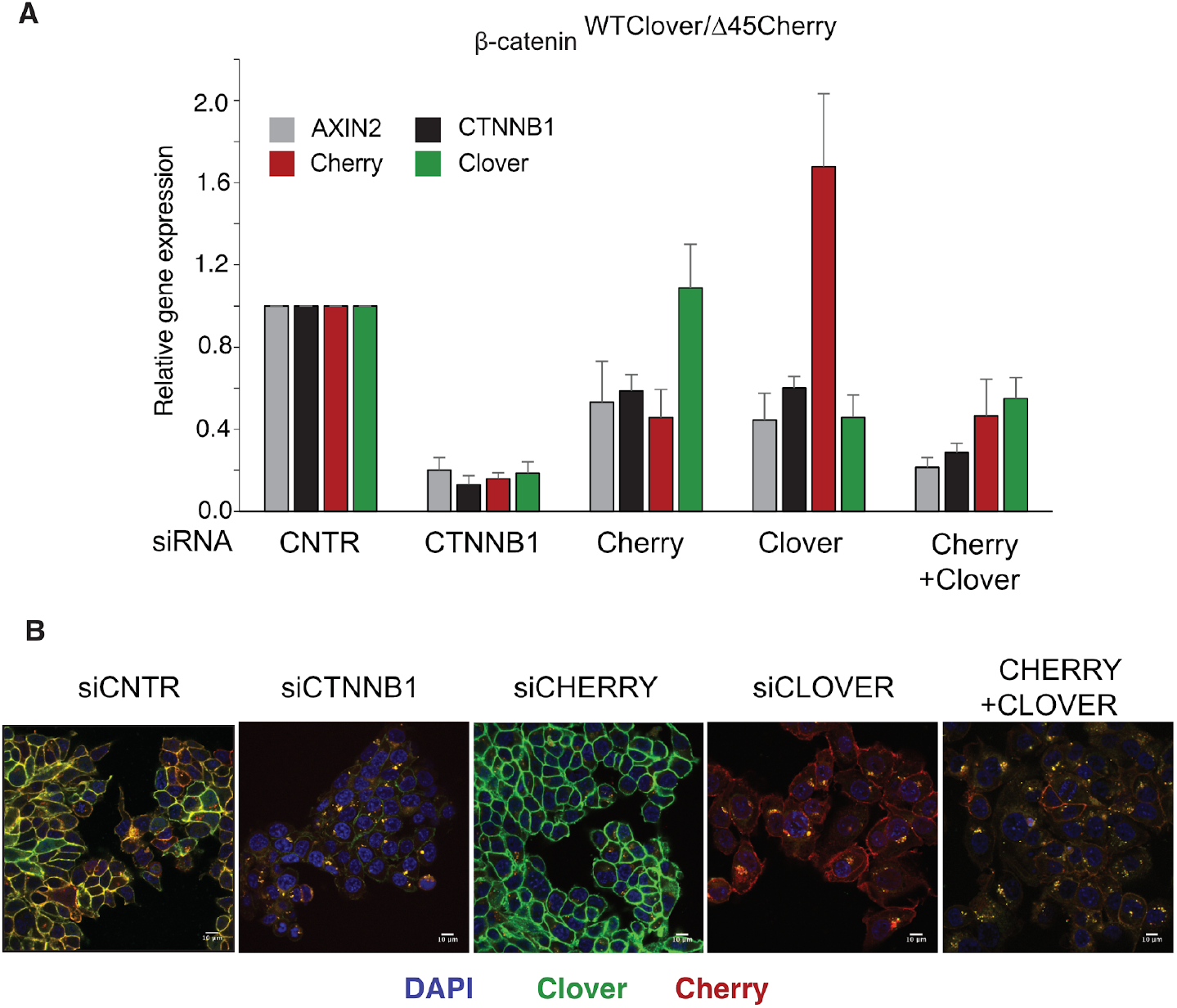
Wild-type and mutant β-catenin both contribute to Wnt pathway activation. (A) Expression levels of *AXIN2, CTNNB1*/β-catenin, *Cherry* and *Clover* were measured 72 hours after knockdown with siRNAs directed against Clover, Cherry or both in HCT116 β-catenin^wtClover/Δ45Cherry^ (n=4, mean ±SD). (B) Immunofluorescence analysis of HCT116 β-catenin^wtClover/Δ45Cherry^ upon transfection with siRNAs targeting CTNNB1, Clover, Cherry or a combination of Clover and Cherry (scale bar: 10μm).

### Fluorescence correlation spectroscopy (FCS) reveals differences in the dynamics of wildtype and mutant β-catenin

FCS is a powerful quantitative microscopy method for measuring local concentrations and diffusional mobilities of fluorescent molecules in cells and tissues, e.g., to determine Wnt ligand-receptor interactions in the cell membrane (Eckert et al., 2020). To simultaneously monitor the dynamics of wild-type Clover-tagged and mutant Cherry-tagged β-catenin of HCT116 cells clone #37, we used FCS with detection in two color channels. With this method, we recorded the fluorescence intensity emanating from a tiny observation volume (ca. 1 fL) as a function of time. For each cell studied, two measurements were performed, with the observation volume first positioned into the cytosol and then into the nucleus. Thus, we focused on the freely diffusing β-catenin fraction, which mediates Wnt signaling, and excluded the membrane-bound fraction, which is responsible for cell-cell adhesion and has been previously shown by FRAP to tightly interact with cadherins, which markedly reduces its mobility (Krieghoff et al., 2006).

From the intensity time traces in the two color channels, autocorrelation functions were calculated and fitted with model functions (Figure S6) to extract two parameters, the diffusional correlation time and the correlation amplitude (Zemanová et al., 2003). The correlation time is inversely proportional to the diffusion coefficient, *D*, of the fluorescent molecules. For this conversion, we performed calibration FCS experiments using fluorescent dye molecules with known *D* values as reference samples (see Methods). According to the Stokes-Einstein law, the diffusion coefficient is inversely related to the linear extension of the diffusing entity and thus to the cube root of its volume/molecular mass. Thus, measurement of *D* values provides size information. The correlation amplitude is inversely proportional to the average number of molecules, *N*, in the observation volume. The calibration experiment with dye molecules mentioned above also allows us to determine the effective volume, *V_eff_*, from which the fluorescence emanates, so that the molecule concentration, *c*, in the observation volume can simply be calculated as *c* = *N* / *V_eff_*. In addition to autocorrelation analysis, we calculated crosscorrelation functions to identify correlated intensity fluctuations in the two color channels. These will appear if the Clover and Cherry labeled β-catenin molecules co-diffuse, i.e., bind to each other and migrate in concert through the observation volume. Cross-correlation amplitudes between wild-type Clover-tagged and mutant Cherry-tagged β-catenin were always found to be zero within the experimental error, implying that these two β-catenin variants diffuse independently and do not associate to any significant degree (Figure S6).

On the basis of FCS experiments in the cytosol of more than 40 cells, we calculated a median *D* value of 10.6 ± 1.8 μm^2^ s^−1^ for mutant Cherry-tagged β-catenin, which is significantly smaller than the one of wild-type Clover-tagged β-catenin (*D* = 17.3 ± 6.1 μm^2^ s^−1^, Figure 6A). The ratio of diffusion coefficients indicates that mutant Cherry-tagged β-catenin diffuses as part of a larger complex, with a more than four-fold larger volume/overall mass than wild-type Clover-tagged β-catenin. For Clover and Cherry overexpressed in HCT116 cells, we measured *D* = 43 ± 8 μm2 s^−1^ in the cytosol. Taking this number and dividing it by the cube root of the molecular mass ratio between wild-type Clover-tagged β-catenin and Clover (120 kDa: 26 kDa), we obtain an estimated *D* = 26 ± 5 μm2 s^−1^ for the tagged β-catenin, which is on the high end of the distribution of *D* values of the respective ensemble (Figure 6A). Thus, Clover-tagged β-catenin may diffuse as an individual entity or as part of a smaller complex (compared to the one that mutant Cherry-tagged β-catenin is bound to).

**Figure 6:**
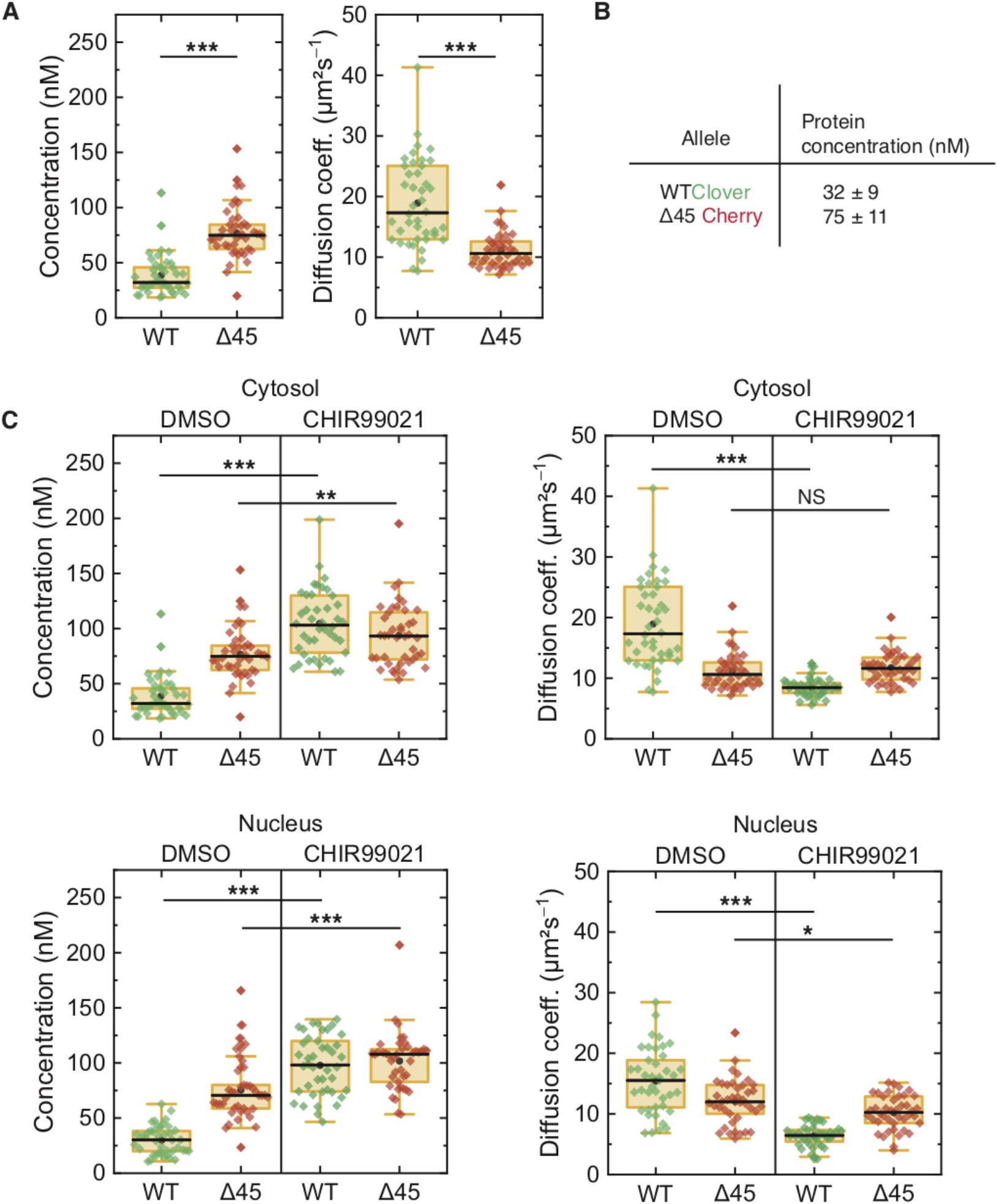
FCS autocorrelation analyses reveal differences in the dynamics and concentrations of wild-type and mutant β-catenin. (A) Concentrations (left) and diffusion coefficients (right) of Clover-tagged wild-type β-catenin and Cherry-tagged mutant β-catenin in the cytosol of HCT116 β-catenin^wtClover/Δ45Cherry^. (B) Protein concentrations (medians of the distributions shown in panel a). (C) Concentrations (left) and diffusion coefficients (right) of Clover-tagged wildtype β-catenin and Cherry-tagged mutant β-catenin measured on HCT116 β-catenin^wtClover/Δ45Cherry^ cells that were treated for 14 – 26 hours with either 10 μM CHIR99901 or DMSO as control. The FCS observation volume was positioned (top) in the cytoplasm and (bottom) in the nucleus. Each data point represents a 120-s FCS measurement in a single cell. In total, more than 40 cells per condition were measured in three independent experiments per box plot. *p* values were calculated with the Mann-Whitney-Test (***< 0.001; **< 0.01; *< 0.05; NS - non-significant). See also Table S1.

From the autocorrelation amplitudes in the green and red color channels, the concentrations of wild-type Clover-tagged β-catenin and mutant Cherry-tagged β-catenin were determined as 32 ± 9 nM and 75 ± 11 nM (medians of distributions over more than 40 cells, Figure 6A and B). These data agree well with previous measurements in the RKO cell line, which revealed β-catenin concentrations of ~52 nM upon Wnt3A stimulation (Hernández et al., 2012).

Next, we were interested whether activation of the Wnt signaling pathway had an impact on the diffusional dynamics of wild-type and mutant β-catenin. Therefore, FCS measurements were performed in the presence of the GSK3β inhibitor CHIR99021, which induces canonical Wnt signaling as depicted in Figure 4A. Under control conditions (DMSO), wild-type β-catenin was less abundant and diffused faster than mutant β-catenin in the cytosol (Figure 6C). Interestingly, treatment with CHIR99021 changed the diffusion coefficients and concentrations of wild-type β-catenin so that they became similar to the ones of mutant β-catenin, whereas smaller effects were observed for the mutant isoform. Essentially, the same effects were obtained for both isoforms in the nuclear fraction (Figure 6C). These observations suggest that, by applying the drug, wild-type β-catenin was also incorporated into a larger complex. The increased concentration suggests that complex formation reduces the probability of β-catenin degradation. Together, these results indicate that GSK3β inhibition renders the concentration and diffusional dynamics of wild-type β-catenin such that they are in the range of those of the mutant isoform.

### APC truncation affects the dynamics of the wild-type β-catenin isoform

*APC* mutations are frequently found in colorectal cancer and define the onset of the transition from adenoma to carcinoma. The complete loss of *APC* is very rare; truncating mutations lacking the C-terminal domain are, however, frequently observed (Cancer Genome Atlas Network, 2012; Fearnhead et al., 2001). Hence, we engineered cell lines carrying mutant truncated APC using CRISPR/Cas9 editing in the biallelically tagged HCT116 clone #37, and quantitatively assessed the impact of APC truncations on wild-type and mutant β-catenin protein in the same cell.

A sgRNA (sgAPCb) was designed by E-CRISP to target the mutation cluster region (MCR) in exon 15 of *APC* (Heigwer et al., 2014; Zhan et al., 2019), introducing a premature stop codon (Figure 7A). Edited single cell clone was verified by amplicon sequencing confirming the homozygous *APC* mutation (Figure 7A). Subsequently, FCS measurements were performed to study the cytosolic and nuclear fractions of both parental HCT116 β-catenin^wtClover/Δ45Cherry^ and HCT116 β-catenin^wtClover/Δ45Cherry^ APC^LOF^ (sgAPC) cells. In the cytosol (Figure 7B) and in the nucleus (Figure S7A) of APC^LOF^ cells, the concentration and diffusional dynamics of wild-type β-catenin were different in comparison to cells with wild-type APC.

**Figure 7:**
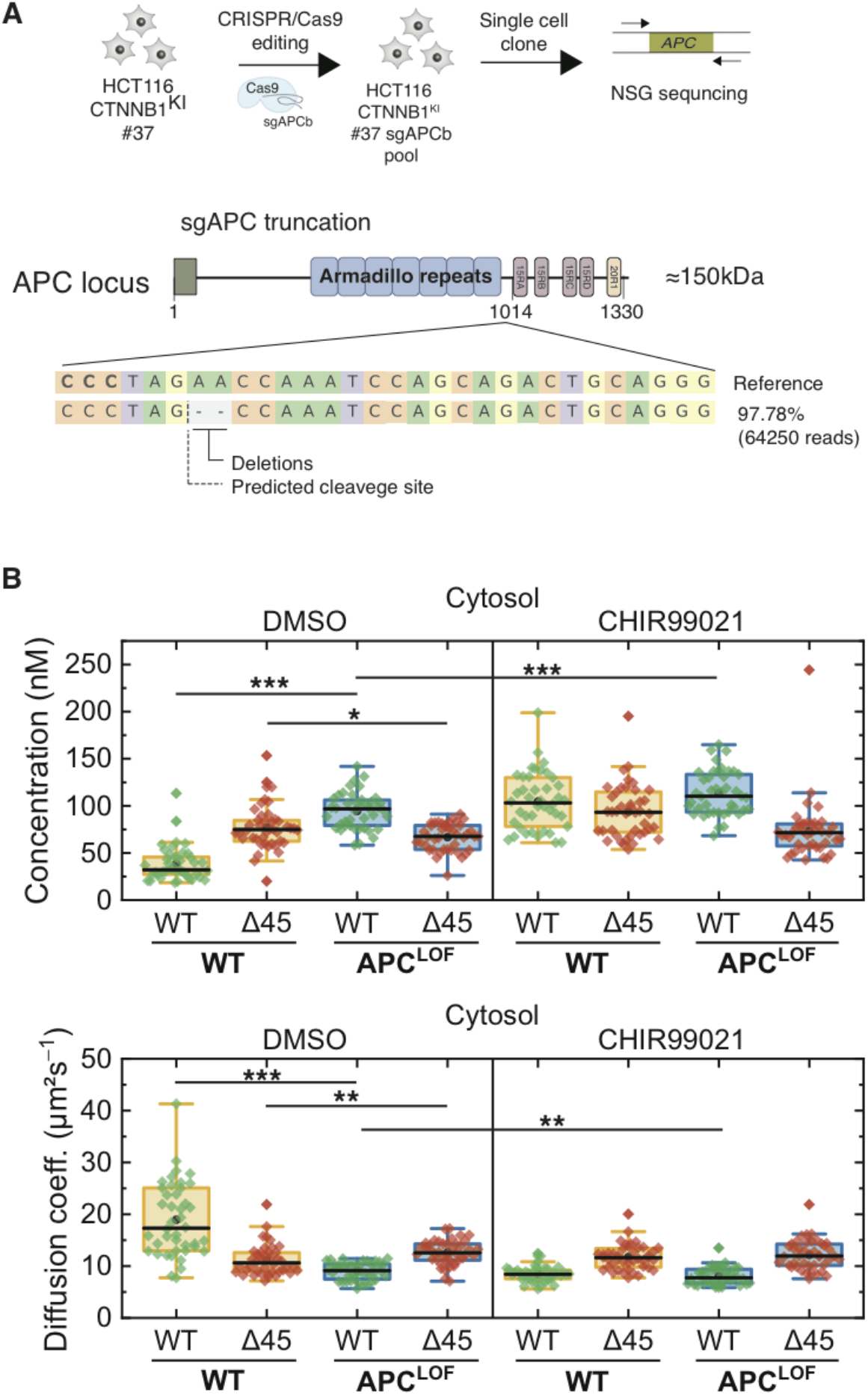
Truncation of APC affects abundance and diffusional dynamics of wild-type but not mutant β-catenin in the cytosol. (A) Schematic representation of the *APC* locus and target site of sgACP in the mutation cluster region (MCR) domain. (B) HCT116 β-catenin^wtClover/Δ45Cherry^ and sgAPC targeted clone (APC^LOF^) cells were treated for ~16 hours with either 10 μM CHIR9901 or DMSO as control. Subsequently, FCS measurements were performed in the cytosol (data shown here) and in the nucleus (data shown in Figure S7). Each data point in the box plots represents a result from a 120-s FCS measurement in a single cell. Per box plot, more than 40 cells were investigated in three independent experiments. *p* values were calculated with the Mann-Whitney-Test (***< 0.001; **< 0.01; *< 0.05; NS - non-significant). The exact values are provided in the Supplementary Table 1.

To investigate whether mutant β-catenin can still be found in the destruction complex, we performed immunoprecipitations with an APC antibody and with GFP/Clover and RFP/Cherry beads. Both wild-type β-catenin-Clover and mutant β-catenin-Cherry bind to APC, Axin1 and GSK3β (Figure S7B) indicating that both isoforms are part of the destruction complex. Both alleles of β-catenin can be regulated by the destruction complex but the level of regulation is different. Mutant β-catenin shows slight regulation upon inhibition of GSK3β or truncation of APC while the wild-type allele is more sensitive to these perturbations.

## DISCUSSION

How disease-causing genetic alleles affect protein function has remained a subject of intense research (Sahni et al., 2015). Mutations in β-catenin that activate the Wnt pathway occur in about 5% of colorectal and more than 25% of liver and uterine tumors (Hoadley et al., 2018; Sanchez-Vega et al., 2018) Nevertheless, the investigation of its biochemical properties has been hampered by the lack of suitable tools to study its function at a physiological level. Here, we generated genome engineered cell lines to simultaneously analyze wild-type and oncogenic mutant alleles of β-catenin. Using CRISPR/Cas9, we generated a bi-allelic endogenously tagged β-catenin cell model carrying mClover-tagged wild-type and mCherry-tagged mutant β-catenin. While tagging of proteins can interfere with protein functions, it can also lead to accumulation of the protein and its potential overactivation. In line with our study previous overexpression studies showed that C-terminal tagging of β-catenin did not affect its functions (Giannini et al., 2000; Jamieson et al., 2011; Kafri et al., 2016; Krieghoff et al., 2006). We also show that all generated clonal cell lines with one or both tagged alleles had the same proliferation capacity (Figure 3A), indicating that the tag does not lead to a loss-of function phenotype. In contrast, inactivation of β-catenin or canonical Wnt signaling in HCT116 cells has been previously shown to inhibit proliferation (Scholer-Dahirel et al., 2011; Voloshanenko et al., 2013). The region required for its adhesion function is within the central part of β-catenin and not at the C-terminus (Valenta et al., 2012; Xing et al., 2008). Therefore, the likelihood of C-terminal tagging affecting this function is rather small. Indeed, we found that β-catenin adhesive function was not affected (Figure 3D,E, S3B,C). Previous studies used overexpression of fluorescently tagged β-catenin to analyze its kinetics (Jamieson et al., 2011; Kafri et al., 2016; Krieghoff et al., 2006) By contrast, here we studied the dynamics of endogenous wild-type and mutant β-catenin by leveraging genome engineering tools for endogenous tagging.

Advances in genome editing methods such as CRISPR/Cas9 enable the modification of endogenous loci and the manipulations of genes for applications such as endogenous tagging of genes (Cong et al., 2013; Dambournet et al., 2014; Mali et al., 2013). Recent reports described several approaches to increase the editing efficiency (Lackner et al., 2015; Martin et al., 2019; Yang et al., 2014). Based on these reports, we designed our two donor templates with 180bp homology arms for the insert of around 3 kbp. Surprisingly, the efficiency of integration of the two donor templates was different, with Clover having about a 12-fold better integration efficiency than Cherry (Figure S1B). This might indicate that the efficiency of HDR depends not only on homology arms but also on the integrated sequence.

Using FCS we analyzed mutant and wild-type proteins in the same cell. These measurements revealed that wild-type β-catenin diffuses faster than the mutant isoform in the cytoplasmic and the nuclear compartments. This difference was abolished when we activated Wnt signaling by inhibition of GSK3β or truncation of APC, resulting in slower diffusion and lower abundance of wild-type β-catenin, similar to the mutant isoform. The changed diffusional dynamics upon activation of canonical Wnt signaling is likely due to the onset of interactions with members of the destruction complex or with transcription factors, respectively. Along this line, FRAP analysis demonstrated that TCF4, APC, Axin and Axin2 reduced β-catenin mobility and nucleo-cytoplasmic shuttling suggesting that these interaction partners control the subcellular localization of β-catenin by sequestering it in the respective compartment (Krieghoff et al., 2006). Accordingly, Goentoro and Kirschner demonstrated that structural changes of β-catenin rather than molecule numbers regulate Wnt signaling (Goentoro and Kirschner, 2009). A recent manuscript demonstrated that wild-type β-catenin resides in slowly diffusing complexes in the cytosol and nucleus in HAP1 cells (de Man et al., 2020), suggesting that Wnt activation regulates β-catenin dynamics by a combination of destruction complex modifications, and modulating nucleoplasmic shuttling and nuclear retention.

β-catenin plays an important role not only in Wnt signaling but also in cell adhesion, in which it links the adhesion complex to the actin cytoskeleton *via* its interaction with the actin-binding protein α-catenin (Rimm et al., 1995). Since the different functions require distinct localization, β-catenin can be found in at least two distinct pools, one at the cell membrane and one that can shuttle between the cytosol and nucleus. Whether both pools originate from a common pool or from two independent pools, and how these reservoirs regulate each other is still a matter of debate. Accordingly, Valenta *et al*. showed *in vivo* that it is possible to affect one function of β-catenin without interfering with the other (Valenta et al., 2012, 2011). Moreover, different molecular forms of β-catenin have been described, demonstrating that most cytosolic β-catenin exists in a monomeric form, whereas dimers of α-catenin and β-catenin interacted mainly with cadherin (Gottardi and Gumbiner, 2004). Interestingly, our immunoprecipitation and FCS analyses indicate that wild-type and mutant β-catenin are found in two separate pools. We observed that β-catenin is mobile in both compartments, whereas mutant β-catenin is found in higher molecular weight complexes.

In summary, we engineered, to our knowledge, the first biallelically tagged cell model carrying a different fluorescent fusion protein on each allele, demonstrating the feasibility of our one-step tagging strategy to analyze the consequences of genetic variants. β-catenin knock-in cell lines provide a powerful tool for investigating β-catenin’s role in the canonical Wnt signaling, and also in adhesion. Due to these properties of β-catenin, these cell models can serve to evaluate both functions in parallel on the endogenous level. Cell models of wild-type and mutant variants can also be used for drug discovery, identifying small molecules that only interfere with mutant, but not wild-type β-catenin, thereby circumventing side effects that are often associated with interfering with homeostatic levels of Wnt signaling. In general, bi-allelic tagging of genetic variants in their endogenous locus can open new avenues towards understanding and possibly targeting disease-causing variants in a broad spectrum of cellular and organismal phenotypes.

## METHODS

### Cell Culture

The parental colon cancer HCT116 (β-catenin^wt/Δ45^) cell line was purchased from the American Type Culture Collection. HCT116 β-catenin^wt+/-^ cells were obtained from Horizon Discovery. Cells were culture in McCoy’s medium (GIBCO) supplemented with 10% fetal bovine serum (Biochrom GmbH). Cells were grown at 37°C and 10% CO_2_ in a humidified atmosphere. Cells were tested for cross-contamination using SNP profiling by a Multiplex human Cell Authentication (MCA) assay (Multiplexion) and regularly checked for Mycoplasma contamination.

### Transfection

Transient transfections were performed using TransIT-LT1 Transfection Reagent (731-0029; VWR) for DNA plasmids and Lipofectamine RNAiMax (Thermo Fisher Scientific) for siRNAs according to the manufacturer’s protocol. 15-25nM Dharmacon siRNAs (Horizon discovery) and 5-10nM Ambion siRNAs (Thermo Fisher) were used for reverse siRNA transfection (Table S2).

### Small molecule inhibitors

GSK-3β inhibitor CHIR99021 (Cat. No. 361571) and Tankyrase inhibitor XAV939 (Cat. No. 575545) were obtained from Merck-Millipore. ICG001 (Cat. No. Cay16257-1) was obtained from Biomol. DMSO was used as the vehicle control.

### Western blot and co-immunoprecipitations assays

Whole cell lysates were extracted using a buffer containing non-ionic detergents such as Triton X-100 or NP-40, supplemented with protease inhibitors (Roche). Protein concentration was determined using BCA protein assay kit (Thermo Fisher Scientific) according to the manufacturer’s instructions. Samples were loaded on 4-12% NuPAGE Bis-Tris gels or 3-8% NuPAGE Tris acetate gels (Thermo Fisher Scientific) and transferred to the nitrocellulose membranes (GE Healthcare, GE10600002) following standard western blotting procedure. Antibodies and their dilutions are listed in Table S3. Clover- and Cherry-tagged proteins were immunoprecipitated by incubation with either GFP-(gta) or RFP-Trap(rta) magnetic beads/agarose (Chromotek) for 1 h at 4°C with rotation. For E-cadherin and β-catenin IPs extracts were incubated with control or respective antibody (Table S3) together with Dynabeads Protein G magnetic beads (Thermo Fisher Scientific) for 14-16 h at 4°C. For signal visualization, blots were incubated with ECL reagent (WBKLS0100; Merck Millipore). Full Western blot scans are provided in the Source Data files Adobe Photoshop was used equally across the whole image for the adjustment of brightness and contrast.

### sgRNA design and cloning

sgRNAs sequences targeting the *CTNNB1* and *APC* genes were designed using E-CRISP (Heigwer et al., 2014), synthesized by Eurofins and cloned into the px459 plasmid (#62988, Addgene) (Table S4), according to a previously described protocol (Mali et al., 2013). To generate donor plasmids the desired homology arm’s regions and epitope tag (either V5 or FLAG) were synthetized by Invitrogen GeneArt Gene Synthesis and cloned into the vector backbone pMK-RQ. The Clover PGK-HygR and Cherry PGK-BRS were cloned from the published vectors pMK278 (Addgene #72794) and pMK282 (Addgene # 72798), respectively (Natsume et al., 2016). PCR primers are listed in Table S6.

### Generation and validation of allele-specific endogenously tagged β-catenin cell lines

HCT116 cells were transfected with px459sgCTNNB1 and the two donor template encoding plasmids KOZAK_FLAG_mClover3 and KOZAK_V5_mCherry using TransIT (731-0029; VWR) according to manufacturer’s instructions. As controls, each donor template was also transfected alone together with px459sgCTNNB1 to generate single-tagged β-catenin cells and to adjust gates for FACS sorting. After 72h, cells were selected with 2 μg/ml puromycin (P9620, Sigma). 48h later, 100μg/ml hygromycin 10687010, Gibco/Thermo Fisher Scientific) and 10μg/ml blasticidin (R21001, Life Technologies GmbH) were added to select edited cells for five days. Subsequently, surviving cells were sorted as single cell clones by FACS into 96 well plates for cultivating. To validate correctly endogenously tagged β-catenin clones genotyping was performed. Genomic DNA was extracted with phenol-chloroform. Different primer pair combinations were designed for genotyping (Table S6). Primer pair (a) binds to the endogenous β-catenin locus flanking the donor cassette indicating homo- or heterozygous integration of the donor template. Primer pairs (b), (b’) and (c) bind outside and inside the donor indicating the correct in-frame integration. Primer pair (d) binds inside the donor indicating which fluorophore was integrated. PCR products were analyzed by agarose gel-electrophoresis.

### TCF4/Wnt-reporter activity assay

To determine Wnt signaling activity, the luciferase-based dual Wnt reporter assay was performed as described previously (Demir et al., 2013). Briefly, cells were transfected with plasmids encoding the TCF4/Wnt-driven firefly luciferase and actin-promoter driven Renilla luciferase as control (Table S4). Dual-luciferase readout was performed 48 hrs after transfection using Mithras LB940 plate reader (Berthold Technologies). Wnt activity was calculated by normalization of the TCF4/Wnt-luciferase values to the actin-Renilla values.

### RT-qPCR

Total RNA from cell pellets was isolated with the RNeasy Mini kit (Qiagen) according to the manufacturer’s protocol. RNA was reverse-transcribed into cDNA using the RevertAid H Minus First Strand cDNA Synthesis kit (Thermo Fisher Scientific). qPCR was performed using the Universal probe library (Roche) with the LightCycler 480 (Roche) instrument according to the manufacturer’s instructions). A list of primers used in this paper for qPCR is shown in Table S5.

### Immunofluorescence assays

Sterile coverslips (Thermo Fisher Scientific) were placed into 12 well plates and cell suspension was added. One day later, cells were treated with the indicated drugs for 20 - 24 h. Cells were fixed using 4% PFA (VWR) for 20 minutes at room temperature. For antibody staining, cells were permeabilized with 0.25% Triton/PBS. After several washing steps, cells were blocked for 1 h with 5% goat serum (v/v) and 5% BSA (v/v) in PBS at room temperature. After overnight incubation with the primary antibody at 4 °C, cells were incubated with fluorescently labeled secondary antibodies (1:250) for 1 h at room temperature in the dark. Antibodies were diluted in PBS with 5% goat serum (v/v) and 5% BSA (v/v). Finally, cover slips were gently transferred onto microscope slides (Carl Roth GmbH) and fixed with Vectashield mounting media containing DAPI solution (Biozol Diagnostica). Fluorescence images were acquired with a Leica TCS SP5 confocal microscope. For live cell imaging, cells were added into μ-slide with a glass bottom (Ibidi) and staining was performed as described. For automated microscopy cells were seeded into 384 well plates and subjected to fixation using the CybiWell Vario robotic system. Fluorescence images were acquired using InCell Analyzer 2200 microscope (GE Healthcare) at 20X magnification in three channels (DAPI, Cy3 and FITC) with four sites per well.

### Image analysis

Image analysis of automatic microscopy was performed using R package EBImage (Pau et al., 2010) and adapted based on previous analysis methods (Carpenter et al., 2006; Fuchs et al., 2010). Nuclei and cell body were segmented and features for intensity, shape and texture were extracted for each cell based on the DAPI, actin and tubulin staining. Features were then summarized per experiment by mean calculation over all cells. As a proxy for cell count, the number of segmented nuclei was used. Images were assembled using Affinity Designer, Biorender and Adobe Illustrator.

### Fluorescence correlation spectroscopy

For FCS measurements, HCT116 cells were cultured without phenol red and seeded in fibronectin (Sigma-Aldrich, St. Louis, MO) coated 8-well Nunc Lab-Tek chambered cover glass (Thermo Fisher Scientific). To this end, fibronectin was diluted at 50 μg/ml in Dulbecco’s phosphate buffered saline (DPBS, no calcium, no magnesium, Sigma Aldrich). Each well of the 8-well chamber was incubated with 200 μL of the solution for at least 1 h at room temperature. Afterwards, the solution was removed by aspiration, and the wells were allowed to dry. Fibronectin coated chambers were either used directly afterwards or stored at 4°C for a maximum of 1–2 weeks and washed with DPBS directly before use. Upon treatment with drugs, the cell culture medium was replaced with a drug-containing medium for ~16 – 26 hours before the measurements. FCS measurements were started ~12 h after seeding the cells at 37°C and with 5% CO_2_.

For FCS analysis, intensity time traces of 120 s duration were acquired in two color channels for 120 s each by using a home-built confocal microscope described previously (Eckert et al., 2020). Clover and Cherry were excited with pulsed 470- and 561-nm laser light of typically 3 μW (1.5 kW/cm^2^) and 5 μW (2.8 kW/cm^2^), respectively. The emission was filtered by 525/50 nm and 607/37 nm (center/width) band pass filters (BrightLine, Semrock, Rochester, NY). For data acquisition, the observation volume of the microscope was first positioned into the cytoplasm and directly afterwards into the nucleus. The intensity decrease during the measurement due to photobleaching was compensated as described previously (Dörlich et al., 2015). Then, autocorrelation and cross-correlation functions were calculated from the time traces, and correlation amplitudes and times were extracted by fitting a model function to the correlation functions that describes free diffusion of fluorescent particles through an observation volume shaped as a three-dimensional Gaussian function (Gao et al., 2017). Two correction factors were applied to the correlation amplitudes of each individual cell measurement to account for photobleaching and uncorrelated background due to cellular autofluorescence. The correction for photobleaching was described recently(Eckert et al., 2020). To correct for autofluorescence according to (Foo et al., 2012), we measured background intensities in the cytosol and in the nucleus for ensembles of more than 15 cells of lines #45 and #33, both under control conditions (DMSO) and with drug treatment (CHIR99021). For each cellular experiment, the correction factor was calculated using the median intensity of the respective ensemble. Finally, the (microscope-dependent) correlation times and corrected amplitudes were converted into (microscope-independent) diffusion coefficients and concentrations, which are reported in this work. To enable this conversion, we carried out careful calibration experiments using solutions of organic dyes (Alexa 488 and Alexa 546, Thermo Fisher Scientific) with well-known diffusion coefficients and concentrations. Furthermore, FCS measurements on fluorescent proteins (eGFP, Clover, Cherry) overexpressed in HCT116 WT cells were performed to obtain estimates of the molecular mass of the β-catenin carrying entities.

## ACKNOWLEDGEMENTS

We are grateful to F. Port, K. E. Boonekamp and S. Redhai for providing critical comments on the manuscripts. We would like to acknowledge all members of the Boutros group and of the SFB1324 for helpful discussions. We thank F. Zhang and MT. Kanemaki for plasmids, which were obtained via Addgene. We are grateful to support by the Excellence Cluster CellNetworks Core Facilities. This work was funded by the Deutsche Forschungsgemeinschaft (DFG, German Research Foundation) Project No. 331351713 – SFB 1324 “Mechanisms and functions of Wnt signaling” (M.B. and G.U.N).

## AUTHOR CONTRIBUTIONS

M.B. and G.A. designed the study; G.A. designed and generated β-catenin alleles in cell lines, characterized and studied different alleles in canonical Wnt signalling; O.V., D.K. characterized endogenous β-catenin alleles; A.E. performed the FCS experiments and analysis; M.B., G.U.N., D.K., O.V., G.A., A.E., wrote the manuscript. M.B. and G.U.N. supervised the study.

## COMPETING INTERESTS

The authors declare no competing financial interests.

## SUPPLEMENTARY TABLES

**Table S1.**
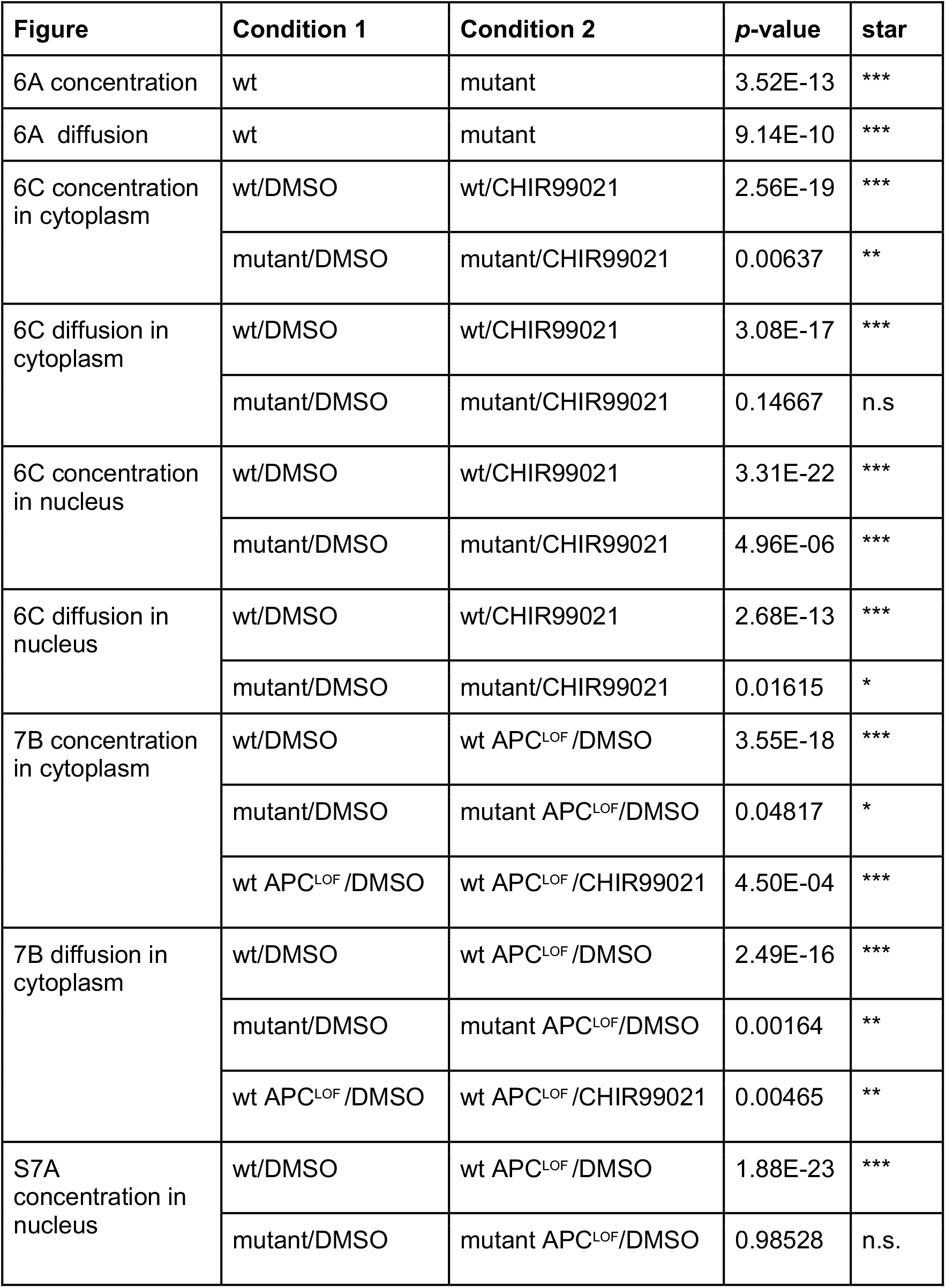
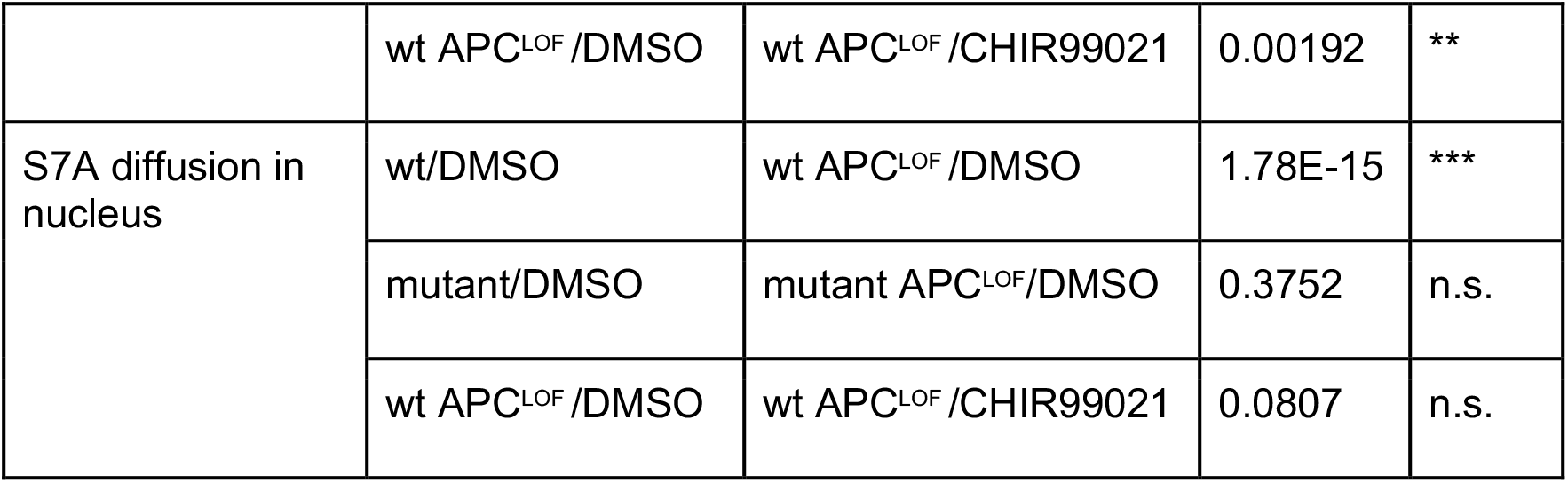

| Figure | Condition 1 | Condition 2 | p-value | star |
| --- | --- | --- | --- | --- |
| 6A concentration | wt | mutant | 3.52E-13 | *** |
| 6A diffusion | wt | mutant | 9.14E-10 | *** |
| 6C concentration in cytoplasm | wt/DMSO | wt/CHIR99021 | 2.56E-19 | *** |
|  | mutant/DMSO | mutant/CHIR99021 | 0.00637 | ** |
| 6C diffusion in cytoplasm | wt/DMSO | wt/CHIR99021 | 3.08E-17 | *** |
|  | mutant/DMSO | mutant/CHIR99021 | 0.14667 | n.s |
| 6C concentration in nucleus | wt/DMSO | wt/CHIR99021 | 3.31E-22 | *** |
|  | mutant/DMSO | mutant/CHIR99021 | 4.96E-06 | *** |
| 6C diffusion in nucleus | wt/DMSO | wt/CHIR99021 | 2.68E-13 | *** |
|  | mutant/DMSO | mutant/CHIR99021 | 0.01615 | * |
| 7B concentration in cytoplasm | wt/DMSO | wt APC<sup>LOF</sup>/DMSO | 3.55E-18 | *** |
|  | mutant/DMSO | mutant APC<sup>LOF</sup>/DMSO | 0.04817 | * |
|  | wt APC<sup>LOF</sup>/DMSO | wt APC<sup>LOF</sup>/CHIR99021 | 4.50E-04 | *** |
| 7B diffusion in cytoplasm | wt/DMSO | wt APC<sup>LOF</sup>/DMSO | 2.49E-16 | *** |
|  | mutant/DMSO | mutant APC<sup>LOF</sup>/DMSO | 0.00164 | ** |
|  | wt APC<sup>LOF</sup>/DMSO | wt APC<sup>LOF</sup>/CHIR99021 | 0.00465 | ** |
| S7A concentration in nucleus | wt/DMSO | wt APC<sup>LOF</sup>/DMSO | 1.88E-23 | *** |
|  | mutant/DMSO | mutant APC<sup>LOF</sup>/DMSO | 0.98528 | n.s. |
|  | wt APC<sup>LOF</sup>/DMSO | wt APC<sup>LOF</sup>/CHIR99021 | 0.00192 | ** |
| S7A diffusion in nucleus | wt/DMSO | wt APC<sup>LOF</sup>/DMSO | 1.78E-15 | *** |
|  | mutant/DMSO | mutant APC<sup>LOF</sup>/DMSO | 0.3752 | n.s. |
|  | wt APC<sup>LOF</sup>/DMSO | wt APC<sup>LOF</sup>/CHIR99021 | 0.0807 | n.s. |

**Table S2.**
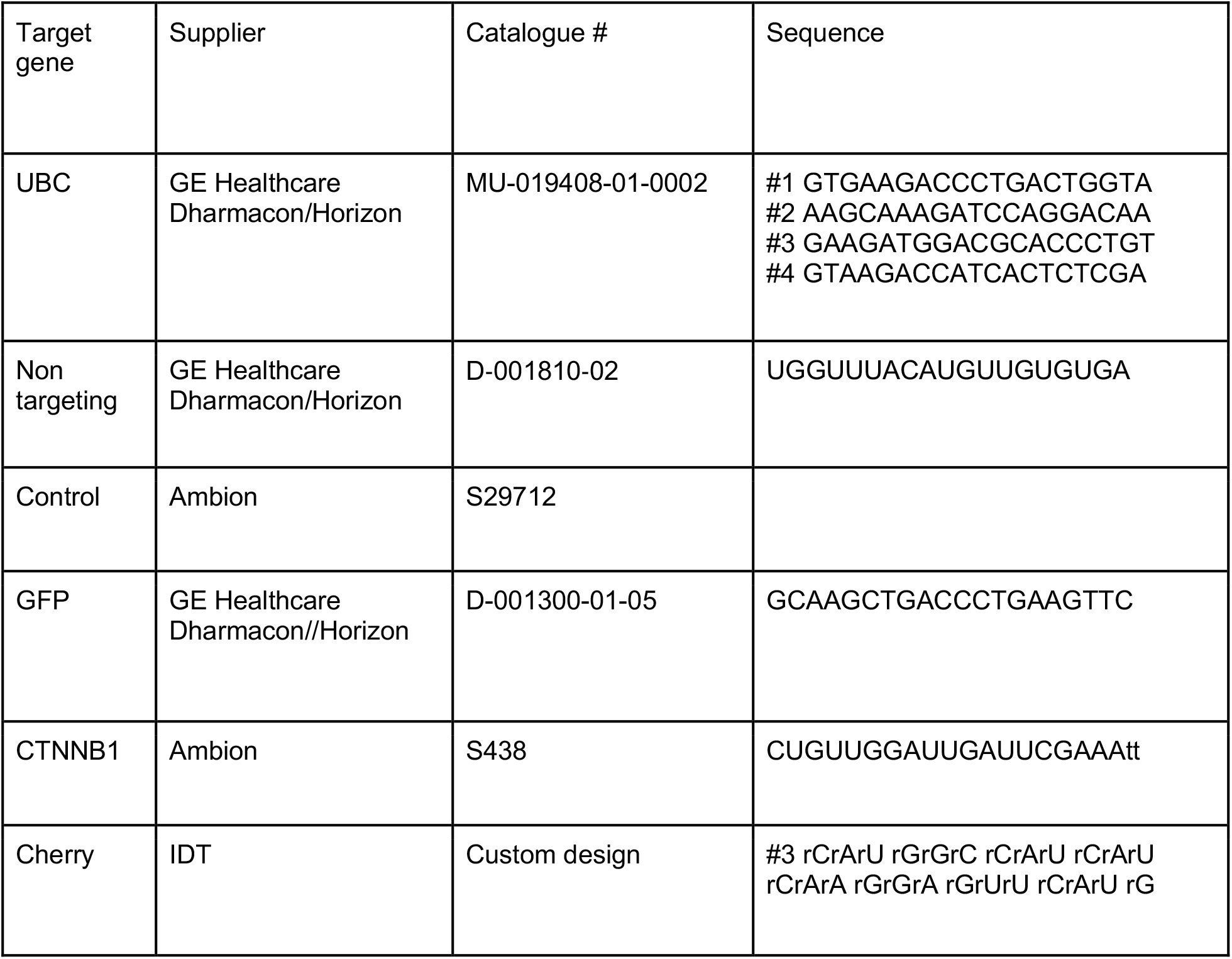
siRNAs

**Table S3.** Antibodies

| Antibody | Company | Catalogue # | Species | Dilution |
| --- | --- | --- | --- | --- |
| β-actin HRP | Santa Cruz Biotechnology | 47778 | rabbit | 1:20000 |
| β-actin | Santa Cruz Biotechnology | 47778 | rabbit | 1:40000 |
| β-catenin | Dianova/Affinity BioReagent | MA1-2001 | mouse | 1:1000 |
| Cherry | Clontech | 632543 | mouse | 1:1000 |
| GFP | Invitrogen | 332600 | mouse | 1:1000 |
| GFP | Invitrogen | A6455 | rabbit | 1:1000 |
| E-Cadherin | BD Biosciences | 610182 | mouse | 1:1000 |
| V5 | Rockland | 600-401-378 | rabbit | 1:2000 |
| V5 | Thermo Scientific | 15253 | mouse | 1:1000 |
| Flag | Sigma | F7425 | rabbit | 1:1000 |
| Flag | Sigma | F3165 | mouse | 1:1000 |
| APC ALI 12-28 | Santa Cruz Biotechnology | sc-53165 | mouse | IP |
| Axin1 C76H11 | Cell Signaling Technology | 2087 | rabbit | 1:1000 |
| GSK3β D5C5Z | Cell Signaling Technology | 12456 | rabbit | 1:1000 |
| Normal IgG | Cell Signaling Technology | 2729 | rabbit |  |
| IgG1 K isotype control | eBioscience | 16-4714-81 | mouse |  |
| Anti-mouse IgG-HRP | Jackson ImmunoResearch | 115-035-003 | Goat | 1:10000 |
| Anti-rabbit IgG-HRP | Jackson ImmunoResearch | 111-035-003 | Goat | 1:10000 |
| True Blot ULTRA Anti-mouse IgG-HRP | eBioscience | 18-8817-33 |  | 1:2500-1:5000 |

**Table S4.**
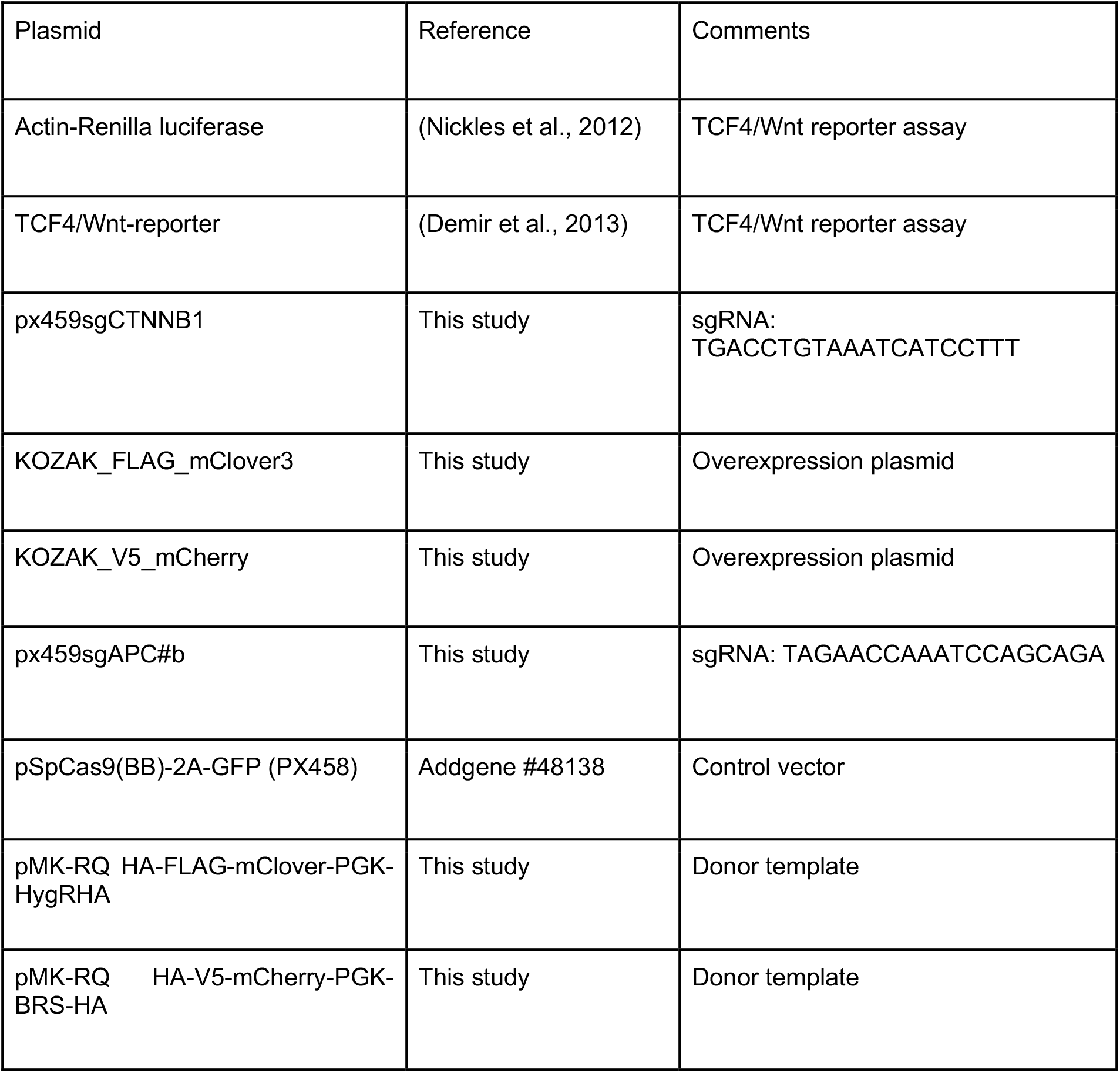
Plasmids

**Table S5.**
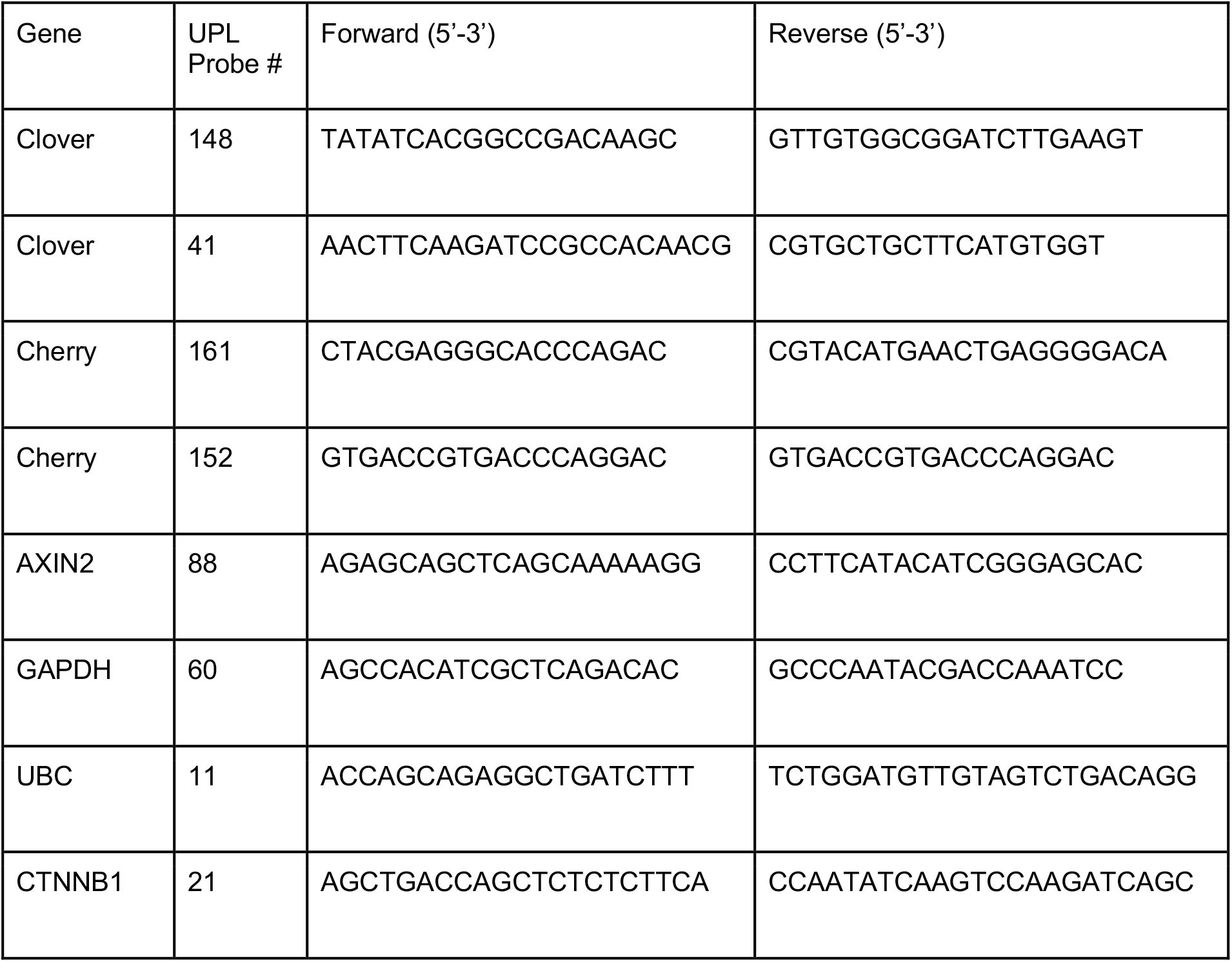
Primers and Roche UPL index used in qPCR experiments

**Table S6.**
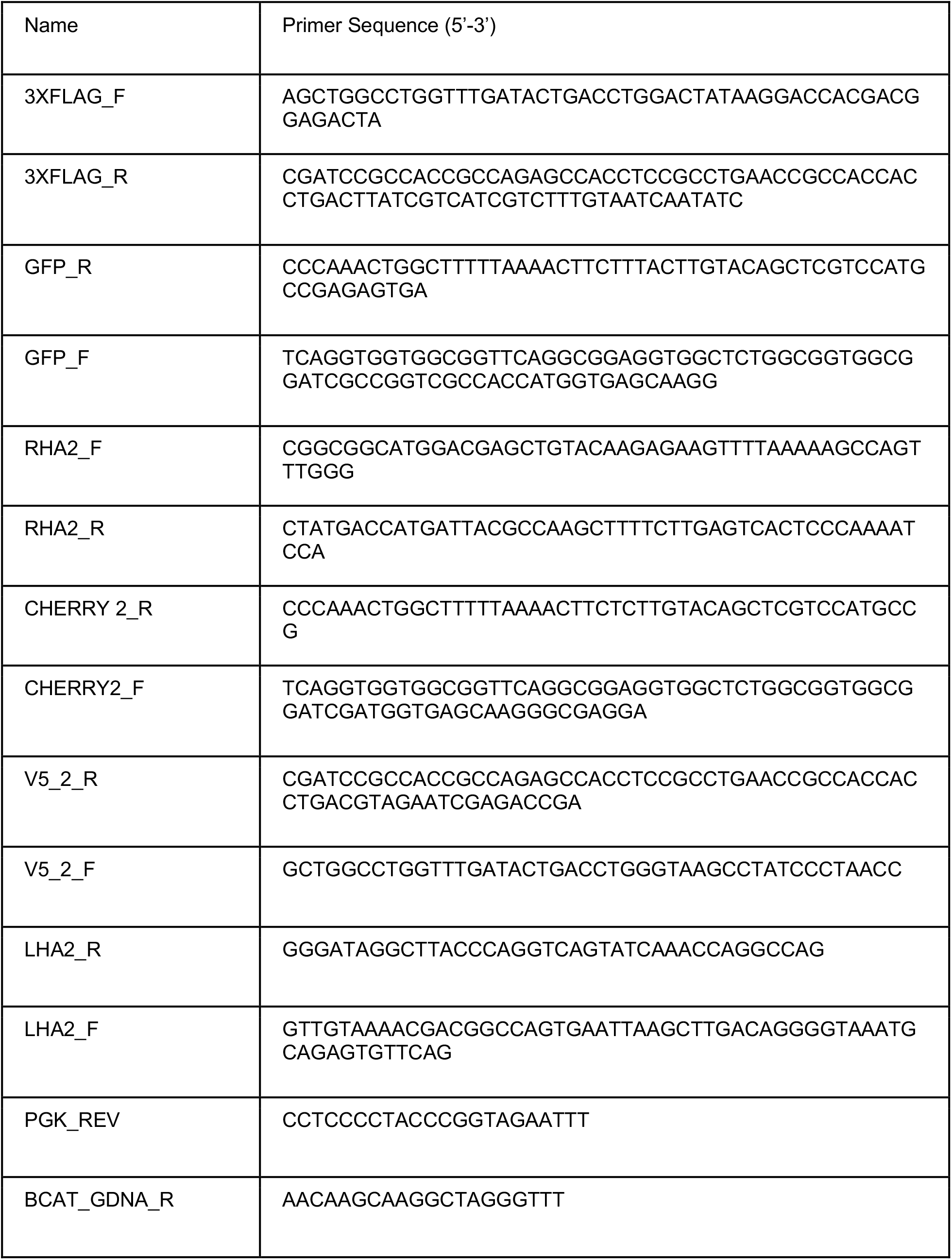
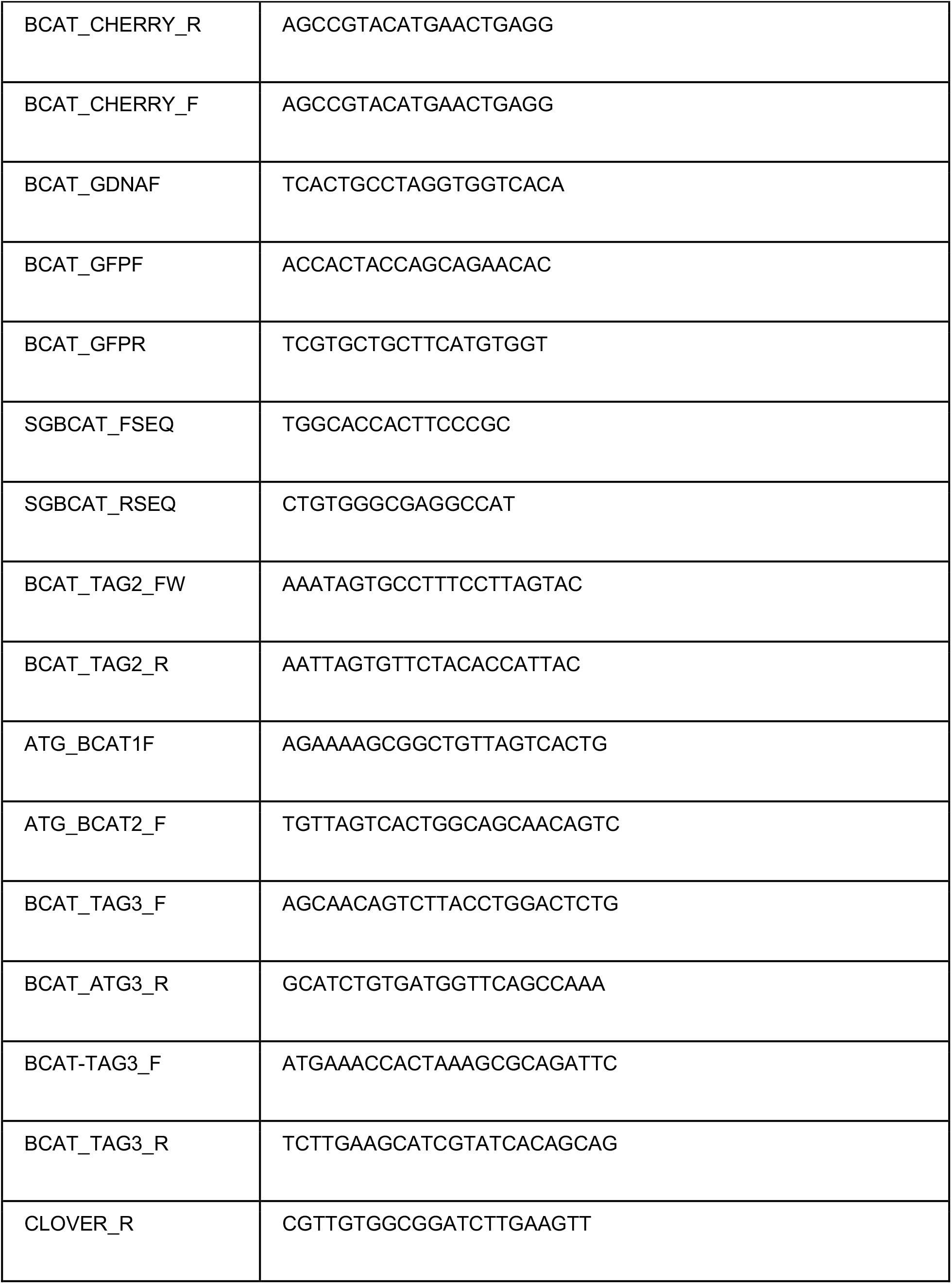
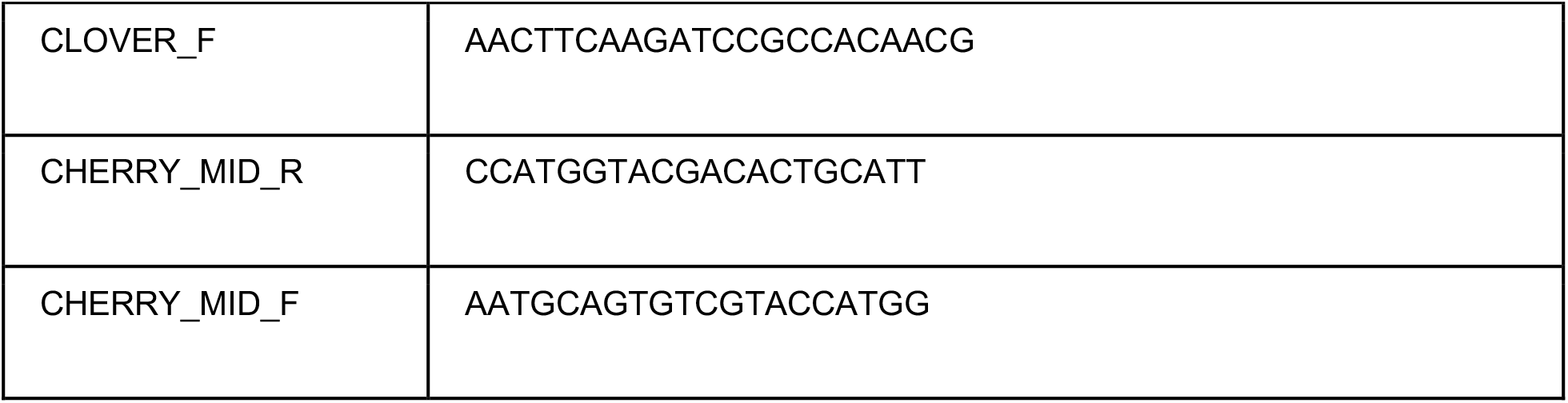
Oligonucleotides

## SUPPLEMENTAL FIGURES

**Figure S1:**
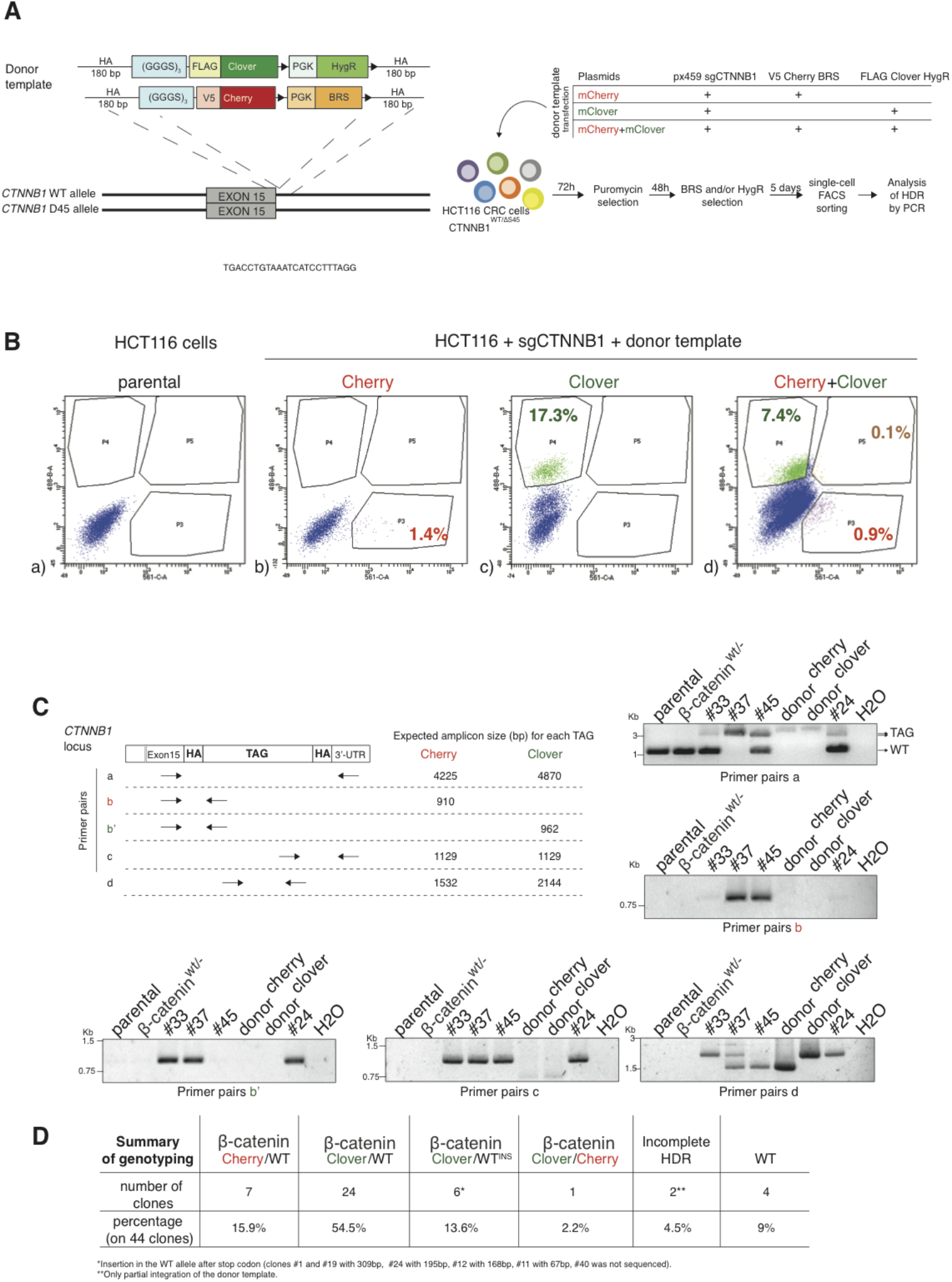
Generation of endogenous fluorescently tagged β-catenin cell lines. (A) Schematic representation of *CTNNB1*/β-catenin tagging strategy and workflow of bi-allelic β-catenin tagged HCT116 cells generation (B) The editing efficiency between the two donor templates was different. The donor template Clover was more efficiently integrated than the Cherry one. (a) Parental HCT116 cells were used as “unstained” sample for FACS analysis (b) HCT116 cells transfected with sgCTNNB1 and Cherry donor template were used to define the gate P3 (mCherry+ cells), (c) HCT116 transfected with sgCTNNB1 and Clover donor template were used to set the gate P4 (mClover+ cells), (d) edited cells transfected with sgCTNNB1 and both Cherry and Clover donor templates are double-positive (gate P5). Percentages of positive cells are shown in each panel. C) The table indicates primers used for genotyping of single cell clones. PCR analysis shows genotyping of HCT116 single cell clones #33, #37, #45 and #24. Parental HCT116 β-catenin^wt/Δ45^, HCT116 β-catenin^wt/-^ and donor templates were used as controls. (D) Table summarizing the obtained HCT116 single clones with their according genotypes.

**Figure S2:**
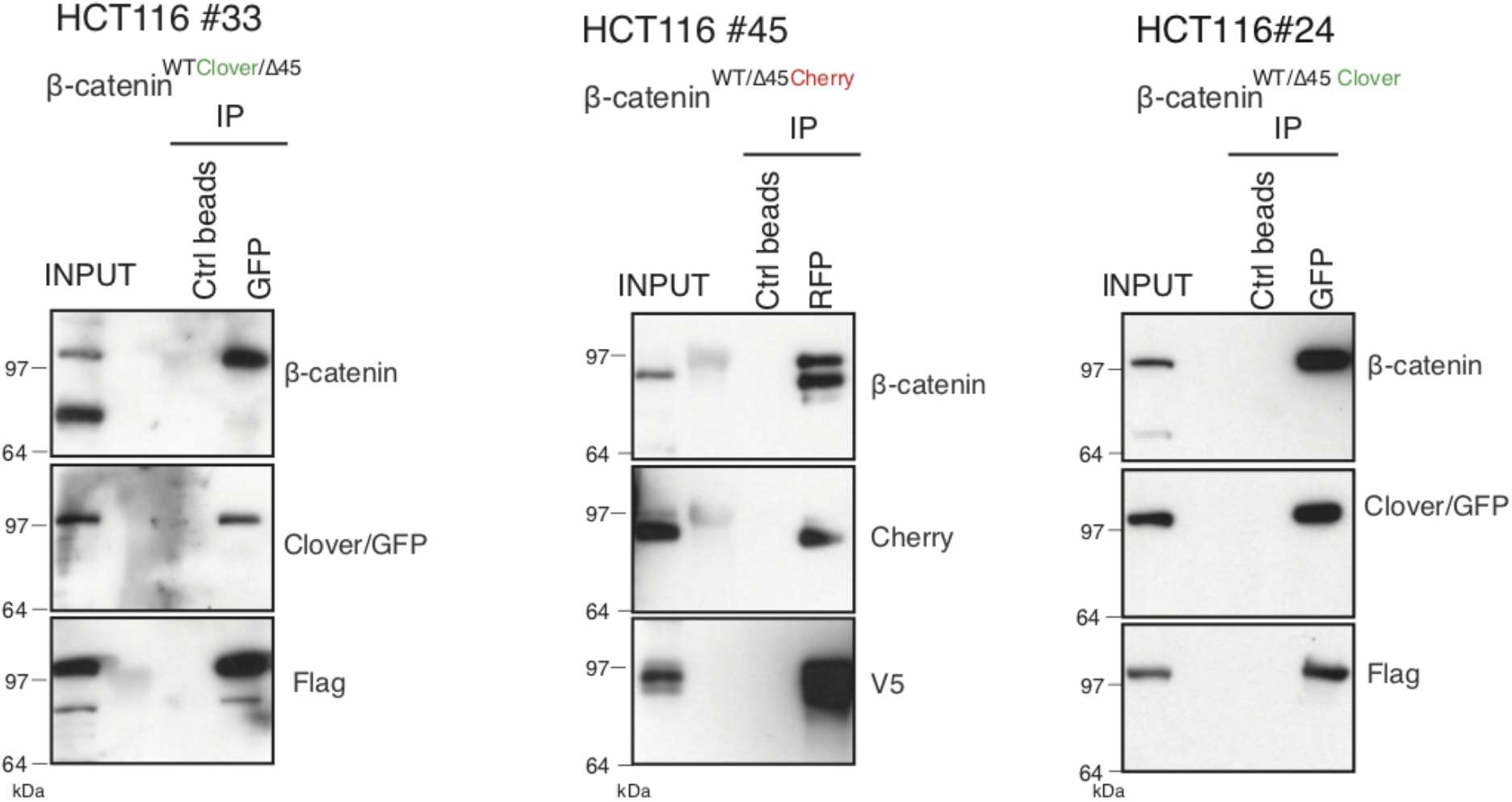
Validation of endogenously fluorescent-tagged β-catenin in HCT116 colon cancer cells. Immunoprecipitation using HCT116 β-catenin^wtClover/Δ45^ (clone #33 - left), β-catenin^wt/Δ45Cherry^ (clone #45 - middle), β-catenin^wtClover/Δ45^ (clone #24 - right) were performed with GFP, Cherry and control beads or with β-catenin antibody, followed by western blotting with the indicated antibodies. Representative results from three independent experiments are shown.

**Figure S3:**
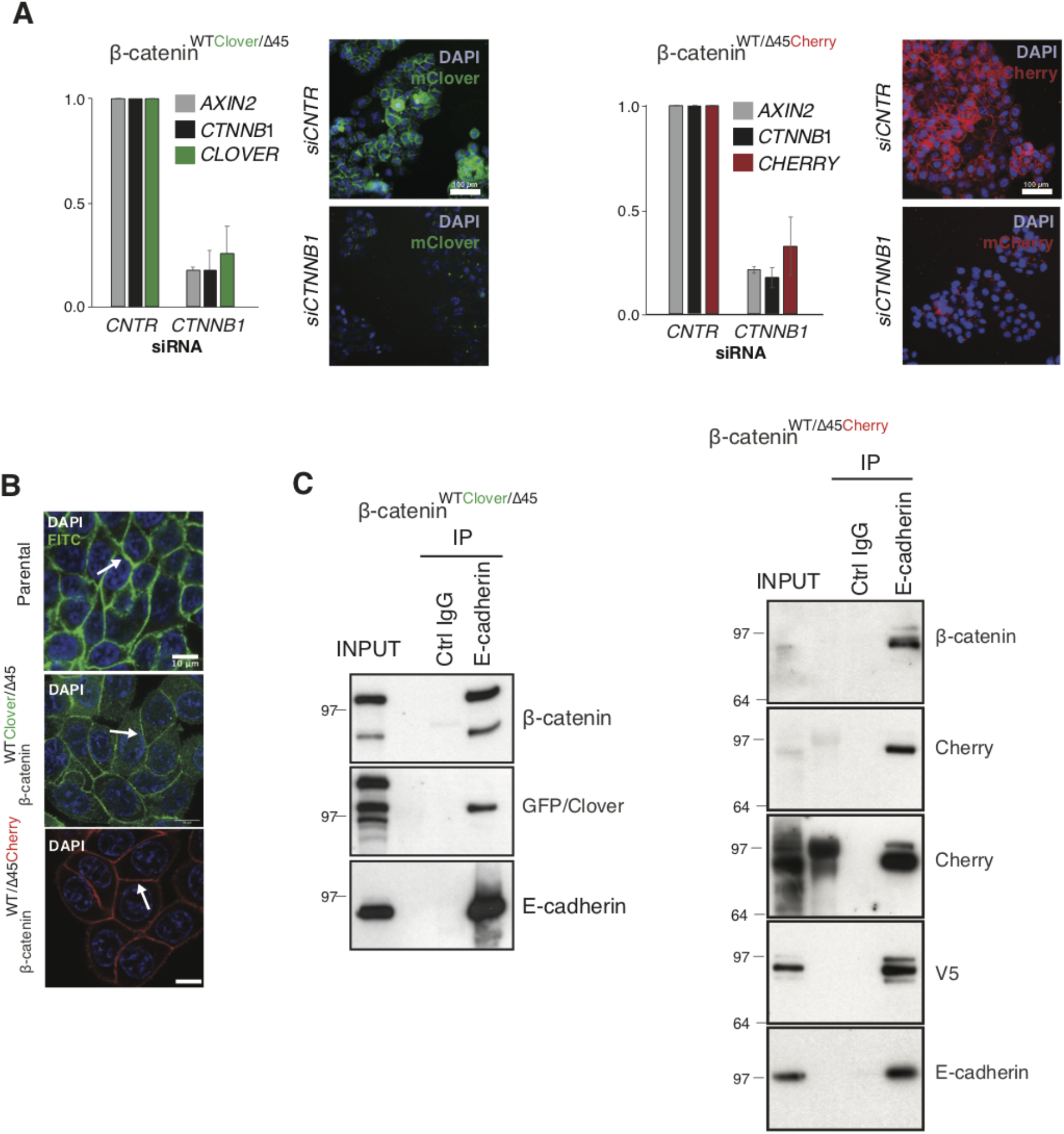
Validation of the physiological function of fluorescently tagged β-catenin. (A) In HCT116 β-catenin^wtClover/Δ45^ and β-catenin^wt/Δ45Cherry^ mRNA-levels of *CTNNB1, AXIN2, CHERRY* and *CLOVER* were determined by RT-qPCR upon silencing of *CTNNB1*/β-catenin. Immunofluorescence analysis of HCT116 β-catenin^wtClover/Δ45^ and β-catenin^wt/Δ45Cherry^ after siRNA-mediated knockdown of *CTNNB1* is shown (n=3; mean ± SEM; scale bar, 100μm). (B) β-catenin accumulates at cell-cell junctions (arrow). Representative immunofluorescence of HCT116 β-catenin^wtClover/Δ45^, β-catenin^wt/Δ45Cherry^ and parental HCT116^wt/Δ45^ stained with β-catenin antibody is shown. Scale bar = 10μm. (C) Immunoprecipitation of HCT116 β-catenin^wtClover/Δ45^ and β-catenin^wt/Δ45Cherry^ with E-cadherin validates its interaction with β-catenin. A representative immunoblot is displayed.

**Figure S4:**
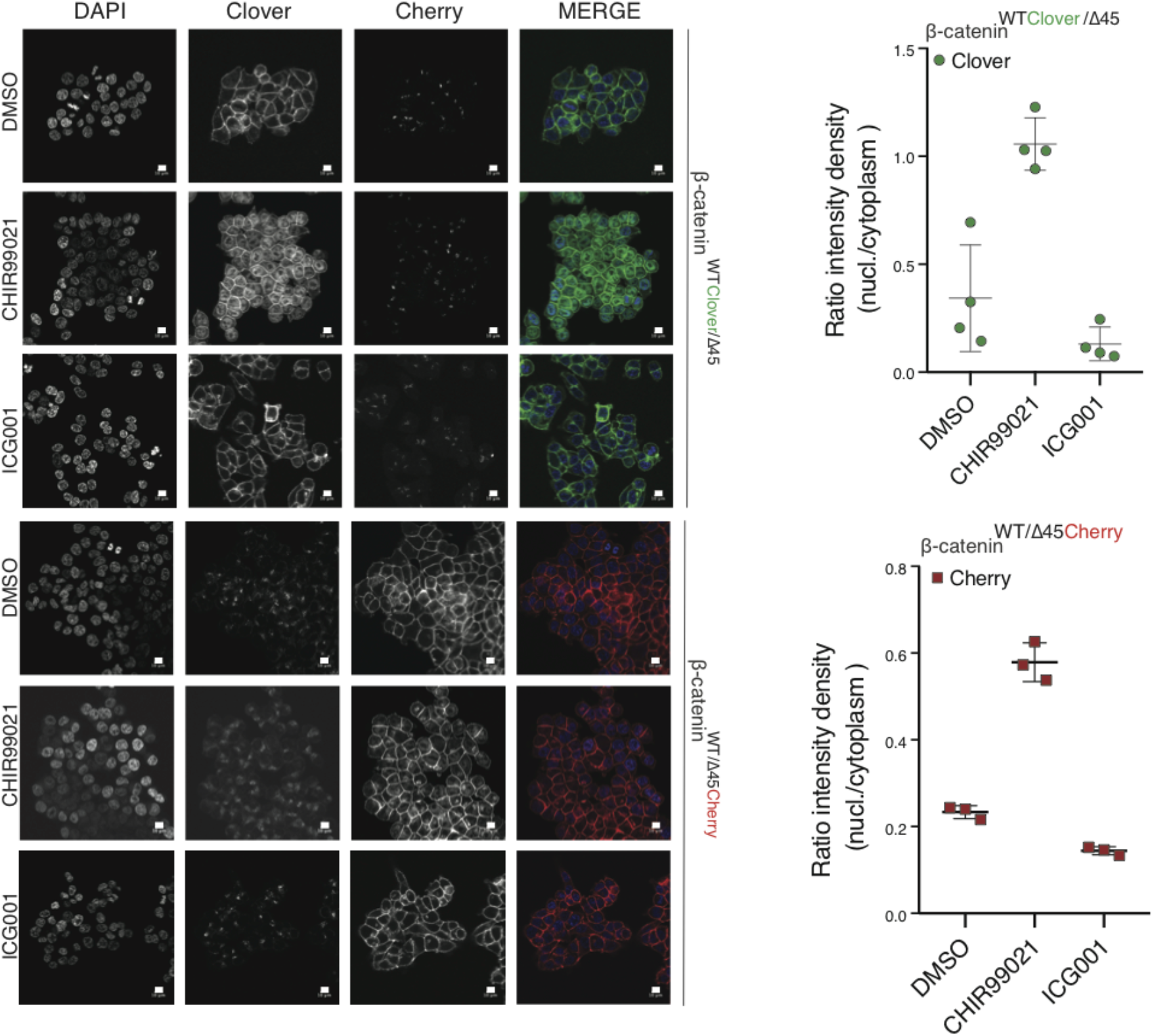
Tagging of β-catenin does not affect its functionality in canonical Wnt signaling. Immunofluorescence analysis of HCT116 β-catenin^wtClover/Δ45^ and β-catenin^wt/Δ45Cherry^ after 24 hours treatment with 10μM CHIR99021 and 10μM ICG-001 is shown. The graphs on the right show the ratio of nuclear to cytoplasmic fluorescence intensities for Clover and Cherry in β-catenin^wtClover/Δ45^ and β-catenin^wt/Δ45Cherry^, respectively (n=3 and 4; mean ± SEM). Scale bar, 10μm.

**Figure S6:**
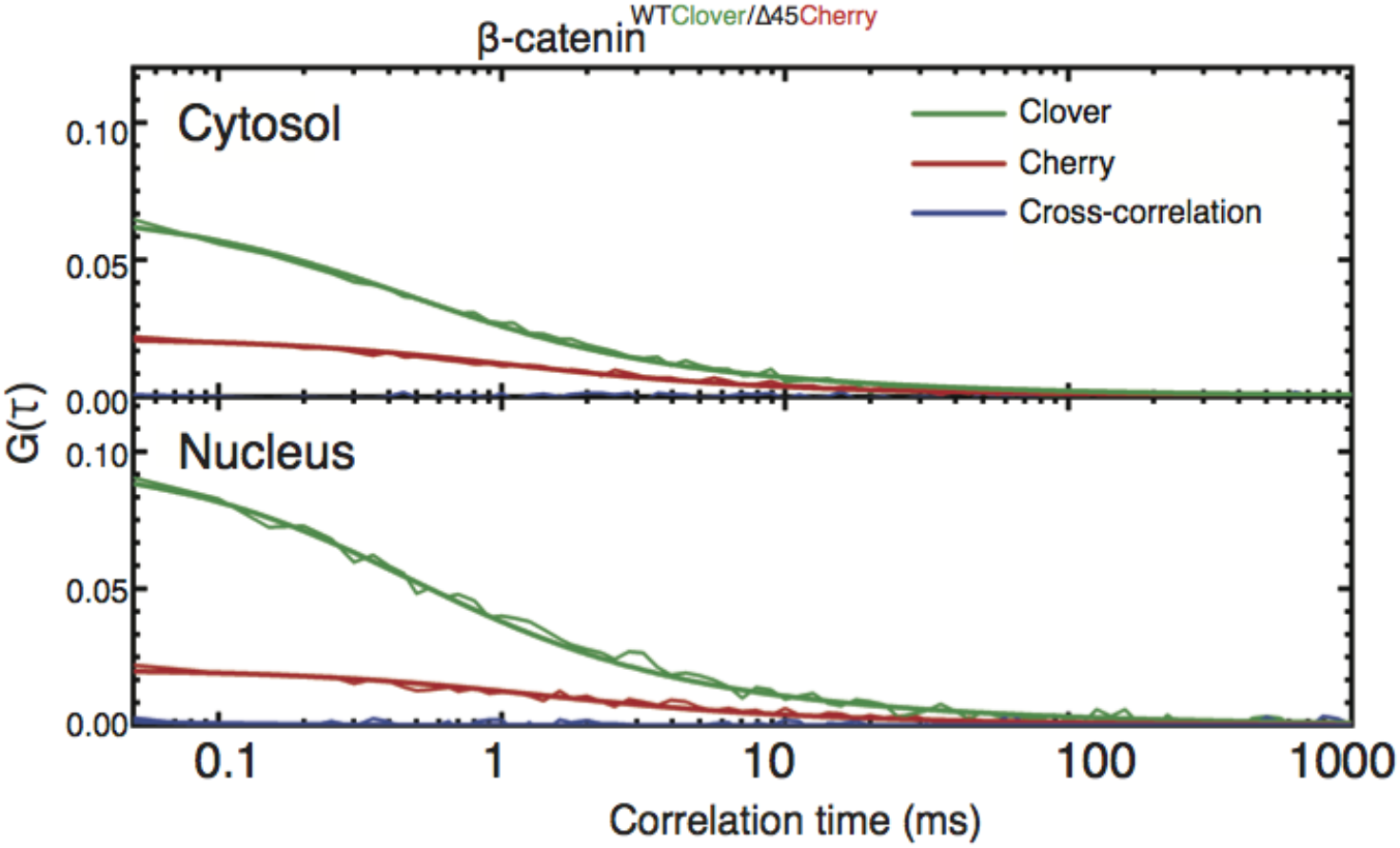
Cross-correlation analysis reveals that the two β-catenin protein isoforms diffuse independently. Shown are representative autocorrelation curves (green, red) and the cross-correlation (blue) from 120-s FCS measurements in the cytosol and the nucleus of HCT116 β-catenin^wtClover/Δ45Cherry^. The non-zero amplitudes of the autocorrelation curves in the green and red color channels indicate the presence of both β-catenin isoforms in the cytosol and in the nucleus. Jagged lines: experimental data, smooth lines, fits with a model function, y-axis: amplitudes of the pair correlation functions.

**Figure S7:**
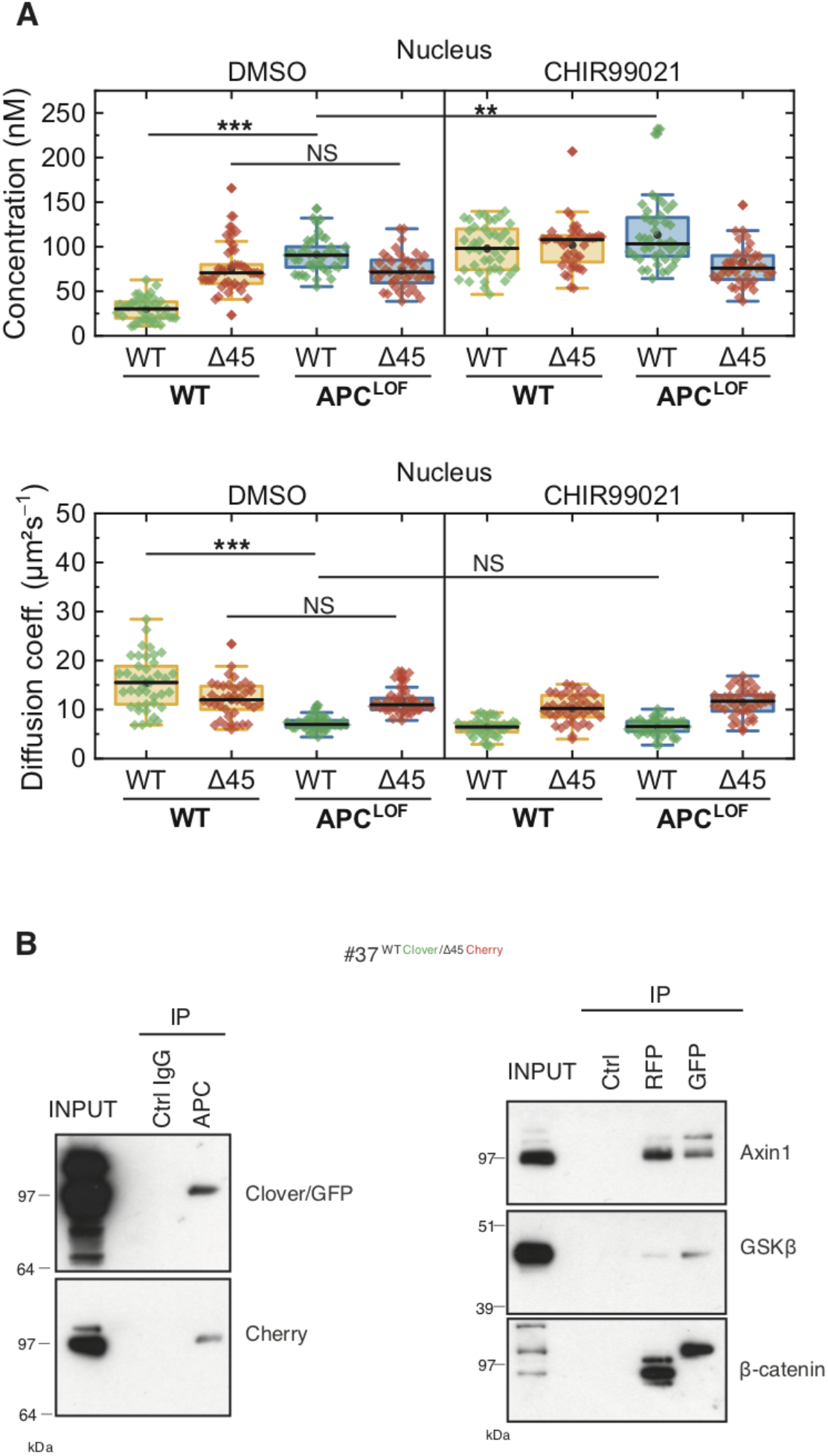
Truncation of APC affects abundance and diffusion of wild-type but not mutant β-catenin in the nucleus. **(A)** HCT116 β-catenin^wtClover/Δ45Cherry^ and sgAPC targeted clone (APC^LOF^) cells were treated for ~16 h with either 10 μM CHIR9901 or DMSO as control. Afterwards, FCS measurements were performed in the cytosol (data shown in Figure 7) and in the nucleus (shown here). For each data point, a 120-s FCS measurement was carried out in a single cell. Per box plot, more than 40 cells were examined in three independent experiments. p values were calculated with the Mann-Whitney-Test (***< 0.001; **< 0.01; *< 0.05; NS - non-significant:) The exact values are provided as the Supplementary Table 1. (B) Both β-catenin alleles of HCT116 cells bind to APC, GSK3β and Axin1. IP with anti-APC antibody was performed GFP/Clover and Cherry were detected (left panel). IPs with RFP/Cherry and GFP/Clover beads were performed and β-catenin, Axin1 and GSK3β were detected (right panel). Representative experiments from 3 independent are shown.

## Notes

### Competing Interest Statement

The authors have declared no competing interest.

